# Temporally integrated single cell RNA sequencing analysis of controlled and natural primary human DENV-1 infections

**DOI:** 10.1101/2020.07.16.206557

**Authors:** Adam T. Waickman, Heather Friberg, Gregory D. Gromowski, Wiriya Rutvisuttinunt, Tao Li, Hayden Siegfried, Kaitlin Victor, Michael K. McCracken, Stefan Fernandez, Anon Srikiatkhachorn, Damon Ellison, Richard G. Jarman, Stephen J. Thomas, Alan L. Rothman, Timothy Endy, Jeffrey R. Currier

## Abstract

Controlled dengue human infection studies present an opportunity to address many longstanding questions in the field of flavivirus biology. However, limited data are available on how the immunological and transcriptional response elicited by an attenuated challenge virus compares to that associated with a wild-type DENV infection. To bridge this knowledge gap, we utilized scRNAseq to analyze PBMC from individuals enrolled in a DENV-1 controlled human challenge study and from individuals experiencing a natural primary DENV-1 infection. While both controlled and natural DENV infection resulted in overlapping patterns of inflammatory gene upregulation, natural DENV infection was accompanied with a more pronounced suppression in gene products associated with protein translation and mitochondrial function, principally in monocytes. This suggests that the immune response elicited by controlled and natural primary DENV infection are similar, but that natural DENV infection has a more pronounced impact on basic cellular processes to induce a multi-layered anti-viral state

## INTRODUCTION

Dengue is one of the most widespread vector-borne viral diseases in the world. The causative agent– dengue virus (DENV) – is a positive-stranded RNA virus maintained in an anthroponotic cycle between the *Aedes aegypti* mosquito and humans (*1*). Consisting of four co-circulating but immunologically and genetically discrete serotypes (DENV-1, −2, −3, and −4), DENV is thought to infect up to 300 million individuals yearly (*2, 3*). Although the majority of DENV infections are subclinical, as many as 100 million infections every year result in symptomatic dengue fever. In its most severe manifestation, dengue fever can progress to dengue hemorrhagic fever/dengue shock syndrome (DHF/DSS) (*4–7*). While the pathogenesis of severe dengue is complex and may involve some degree of genetic predisposition, severe symptoms are more likely to occur in individuals previously infected with a heterologous viral serotype compared to individuals without any preexisting DENV immunity. Despite decades of study, the precise mechanisms underpinning this unique epidemiological feature of DENV infection remain unresolved and continues to impede the development of an effective DENV vaccine.

In addition to the unclear role of preexisting immunity and other host/environmental factors on the immunopathogenesis of DENV infection, the fundamental immunological and molecular signatures associated with early DENV infection remain mostly unresolved. This is partially attributable to the endemic nature of DENV – resulting in frequent and unpredictable exposures in susceptible individuals - as well as the wide variation in the kinetics of asymptomatic incubation/infection prior to any symptomatic manifestation of infection (*8*). Resolving the early unique immunological and molecular signatures specific to DENV infection would aid in the development of rapid diagnostic tools, as well as provide insight into the basic pathophysiology of infection and disease.

DENV human infection models (DHIMs) offer a unique opportunity to formally address many of the outstanding questions in the field of DENV biology. By exposing pre-screened individuals to DENV in a controlled setting, it is possible to closely monitor the immunological response elicited by infection prior to the onset of any traditional clinical symptoms. In addition to providing a platform to answer many fundamental questions of flavivirus biology, DHIMs are a powerful tool to systematically develop and down-select candidate vaccine platforms or therapeutic agents in a controlled setting before expanding to large-scale efficacy trials. Indeed, Kirkpatrick and colleagues have demonstrated the utility of this approach by utilizing an attenuated DENV-2 challenge virus to test the efficacy of the TV003 DENV vaccine product prior to the initiation of larger efficacy trials (*9*).

While the earliest DHIM studies performed in the 1930s utilized unattenuated wild-type DENV strains, all current DHIM studies utilize highly characterized and attenuated viral strains. Viral attenuation is critical to ensure an acceptable safety profile, but it leaves open the possibility that the causative mutations may significantly alter the immunological response elicited by infection, thereby reducing the utility of the model. Limited data are available on how closely these controlled DENV infections mimic the immunological subtleties and complexities of natural primary DENV infection. The majority of DHIMs appear to recapitulate the basic clinical features associated with mild natural primary DENV infection, including rash, myalgia, headache, and fever, as well as the crude virologic/serologic features of infection such as peripheral viremia/RNAemia and DENV-specific seroconversion (*10, 11*). However, no study to date has attempted to directly compare/contrast the molecular signatures associated with either natural or experimental primary DENV infection with any degree of cellular and/or temporal resolution. Understanding how closely a DHIM mimics the molecular subtleties of a wild-type DENV infection would inform decisions on how best to utilize the tool for product down-selection, as well as provide insight into the basic molecular response elicited by DENV infection with an unparalleled degree of temporal resolution.

To close this significant knowledge gap, we utilized high-throughput single-cell RNA sequencing (scRNAseq) technology to assess the longitudinal transcriptional profile associated with both controlled and wild-type primary DENV-1 infection with single-cell resolution. This study utilized samples collected as part WRAIR/SUNY phase one open label DENV-1 human challenge study performed in Syracuse, NY (*10*), as well as samples collected from children enrolled in a hospital-based acute dengue study in Bangkok, Thailand (*12*). Unfractionated PBMC from 8 time points (days 0, 2, 4, 6, 8, 10, 14/15 and 28) from 3 individuals enrolled in the DHIM study were analyzed, as well as 3 time points (acute 1, acute 2, day 180) from two individuals experiencing a natural primary DENV-1 infection. This temporally integrated analysis resulted in the capture of 171,208 cells, which upon examination collapsed into 22 statistically distinct populations corresponding to all major anticipated leukocyte subsets. While all annotated cell populations demonstrated significant and consistent perturbations in their transcriptional profile in response to either natural or experimental primary DENV infection, conventional monocytes respond most robustly to infection across all subjects and study groups from an unbiased transcriptional perspective. Using these data, conserved Differentially Expressed Genes (cDEGs) induced or suppressed by natural or experimental primary DENV were identified, and the overlap between the two arms of the study assessed. The infection-induced cDEGs associated with experimental DENV infection were found to reflect a subset within the larger gene set associated with natural primary DENV infection, primarily corresponding to gene products associated with a cellular response to systemic inflammation and interferon (IFN) production. In contrast, the number infection-suppressed cDEGs was higher in cells obtained following natural primary DENV infection than in cells obtained following experimental DENV infection. Infection-suppressed cDEGs primarily corresponded to gene products associated with protein translation/elongation and mitochondrial function, two cellular processes known to be suppressed by prolonged IFN signaling. These results are consistent with the concept that the immune response elicited by DHIM represents a tempered version of that generated in response to a natural DENV infection, and that the more pronounced inflammation associated with natural primary DENV infection has a correspondingly more pronounced impact on basic cellular processes to induce a multi-layered/systemic anti-viral state. These data provide insight into the molecular level response to DENV infection, and how viral pathogenesis correlates with immune activation and cellular pathophysiology.

## RESULTS

### Subject selection and characterization

The primary objective of this study was to determine the kinetic transcriptional signature associated with experimental DENV-1 infection and to determine how closely this profile correlates with the transcriptional signature accompanying natural primary DENV-1 infection. To this end, three subjects from the SUNY/WRAIR DENV-1 DHIM study were selected for analysis (**Figure 1A**) (*10*). All subjects included in this study received 3.25 × 10^3^ PFU of the 45AZ5 DENV-1 challenge virus strain following extensive pre-screening to ensure an absence of preexisting DENV immunity. All three subjects exhibited significant DENV-1 RNAema between 5 and 15 days post-challenge and demonstrated classic staggered IgM/IgG seroconversion between study days 13 and 16 (**Figure 1A**). PBMC from a total of 8 time points per subject were selected for this study, corresponding to study days 0, 2, 4, 6, 8, 10, 14/15, and 28 (**Figure 1B**). In addition, PBMC from two children experiencing serologically-confirmed natural primary DENV-1 infections were selected for analysis (**Figure 1B, Supplemental Figure 1, Supplemental Table 1**). A total of 3 time points per subject were analyzed: two sequential samples collected during the acute (febrile) phase of infection, and a reference sample collected 6 months post-defervescence. For both the DHIM and natural primary DENV-1 infection subjects, each sample was analyzed in parallel by scRNAseq with TCR/BCR recovery and by flow cytometry (**Figure 1B, Supplemental Figure 2, Supplemental Figure 3**).

**Figure 1.**
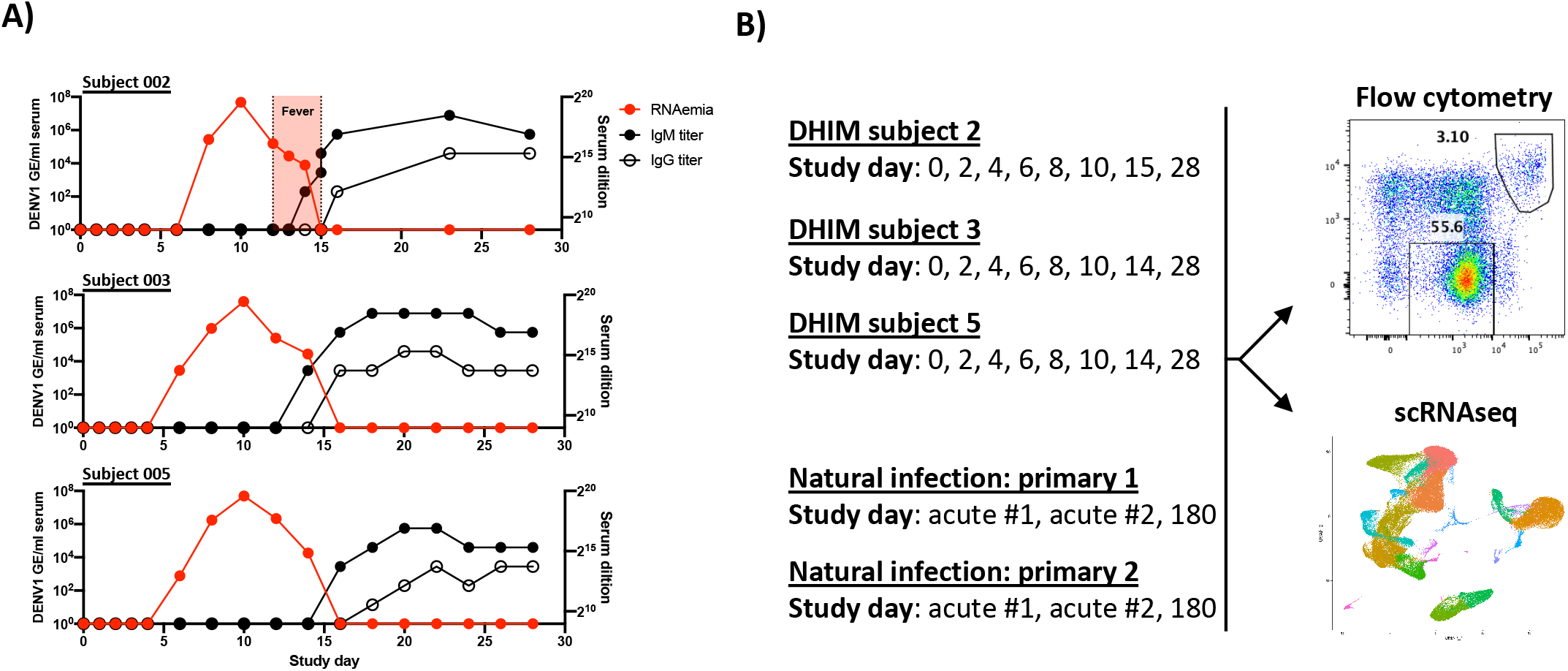
Subject characterization and experimental overview. **A)** Kinetics of RNAemia and DENV IgM/IgG seroconversion in DHIM subjects **B)** Schematic representation of the analysis scheme utilized in this study

### Leukocyte population identification and quantification

In order to assess the transcriptional signature associated with experimental and natural primary DENV infection, high-throughput scRNAseq analysis was performed using the 10xGenomics 5’ capture gene expression platform with both TCR and BCR recovery. Samples were sequenced to achieve an average depth of 108,000 reads per cell, with an average of 5,707 cells captured per library (**Supplemental Table 2**). This final dataset contains a total of 171,208 high quality cells and 22 statistically distinct populations corresponding to all major anticipated leukocyte subsets (**Figure 2A, Figure 2B, Supplemental Table 3, Supplemental Table 4**). With the exception of neutrophils, all annotated populations were consistently observed in all DHIM and natural primary infection samples included in this analysis (**Figure 2C, Supplemental Table 3, Supplemental Table 4**). The relative distribution of all major leukocyte populations was consistent within each subject across all analyzed time point, with the notable exception of activated T cells and plasmablast phenotype B cells, both of which expanded in response to infection in the DHIM subjects on study days 14/15 (**Figure 2D**). No T cell or B cell activation/expansion was observed in the natural primary DENV infection samples, constant with the time points analyzed (*13*). To validate the population annotations as defined by scRNAseq, conventional flow cytometry was performed in parallel on the same samples and the frequencies of major leukocyte populations assessed (**Supplemental Figure 2, Supplemental Figure 3**). The frequencies of all major leukocyte populations captured by either scRNAseq or flow cytometry exhibited a high degree of correlation across all time points in the DHIM sample set (**Supplemental Figure 4**). No cell-associated DENV transcripts were observed in the dataset despite the presence of RNAemia in all subjects.

**Figure 2.**
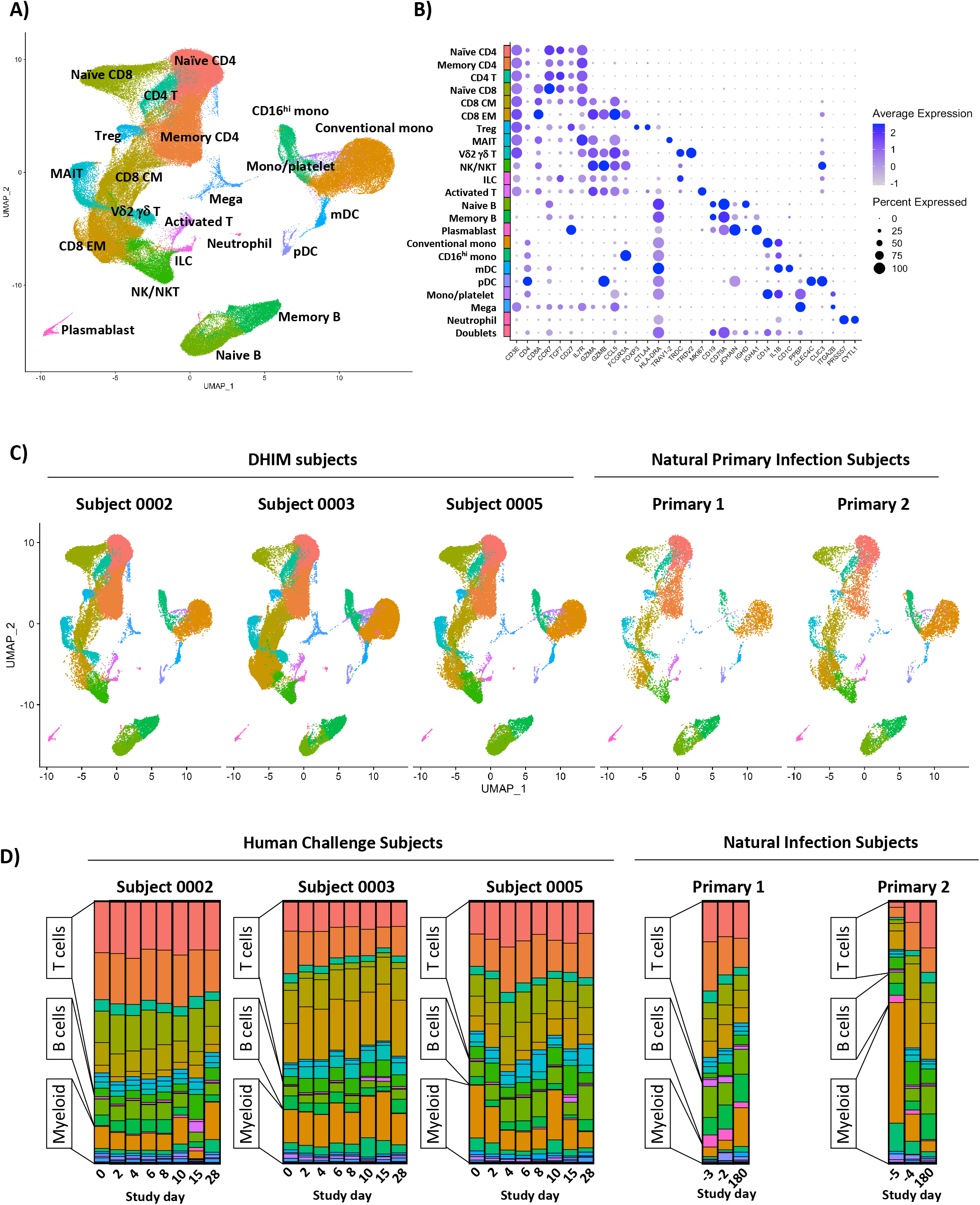
Annotation and quantification of major leukocyte populations following experimental or natural primary DENV infection. **A)** Integrated UMAP projection of scRNAseq data from all annotated leukocyte populations derived from all DHIM subjects (n =3, 8 time points per subject) and natural primary DENV infection subjects (n = 2, 3 time points per subject). **B)** Expression of key linage defining gene products across all annotated leukocyte populations captured in this analysis. **C)** Integrated UMAP projection of all annotated leukocyte populations split by subject. **D)** Relative population abundance in all subjects split by time point.

### DHIM-associated lymphocyte expansion

Three statistically distinct population of B cells were identified within the aggregated DHIM scRNAseq dataset (**Figure 3A**). Based on the expression of canonical linage-associated gene products, these populations were annotated as Naïve B cells, Memory B cells, and plasmablasts (**Figure 3A, 3B**). While the relative frequency of naïve B cells and memory B cells remained fairly consistent throughout the DHIM time course (**Figure 2D, Supplemental Table 3**), robust plasmablast expansion was observed in all subjects on DHIM study days 14/15 (**Figure 3C, Supplemental Figure 4**). Analysis of the full length immunoglobulin sequences associated with these cells revealed that naïve-phenotype B cells expressed IgD or IgM class-switched antibodies with very low levels of somatic hypermutation (SHM), while memory-phenotype B cells expressed an assortment of IgA, IgG and IgM class-switched antibodies with significantly higher overall SHM burdens (**Figure 3D**). Consistent with what has been described following natural primary DENV-1 infection (*14*), plasmablast-phenotype B cells captured on DHIM study days 14/15 expressed a nearly equal mix of IgA, IgG, and IgM class-switched antibodies with relatively high SHM burdens as assessed by both scRNAseq and flow cytometry (**Figure 3D, Supplemental Figure 5**). To confirm the antigen specificity of these putative DENV-elicited plasmablasts, 15 monoclonal antibodies (mAbs) were synthesized from DHIM subject 0002 (5 IgM isotype, 5 IgG isotype, 5 IgA isotype) and tested for their ability to bind DENV-1, −2, −3, and −4 (**Supplemental Table 5**). Consistent with previously published reports describing the antigen specificity of natural or experimental primary DENV infection (*14, 15*), 2 of 15 (13%) of the tested mAbs exhibited DENV-binding activity, including serotype specific and cross-reactive profiles (**Supplemental Table 6**). While the overall magnitude of plasmablast expansion observed following DHIM-1 challenge is lower than what has been described in natural primary DENV-1 infection (*13, 14, 16, 17*), these data suggest that the B cell activation profile and antigen specificity associated with DHIM mirrors that observed in natural primary DENV-1 infection.

**Figure 3.**
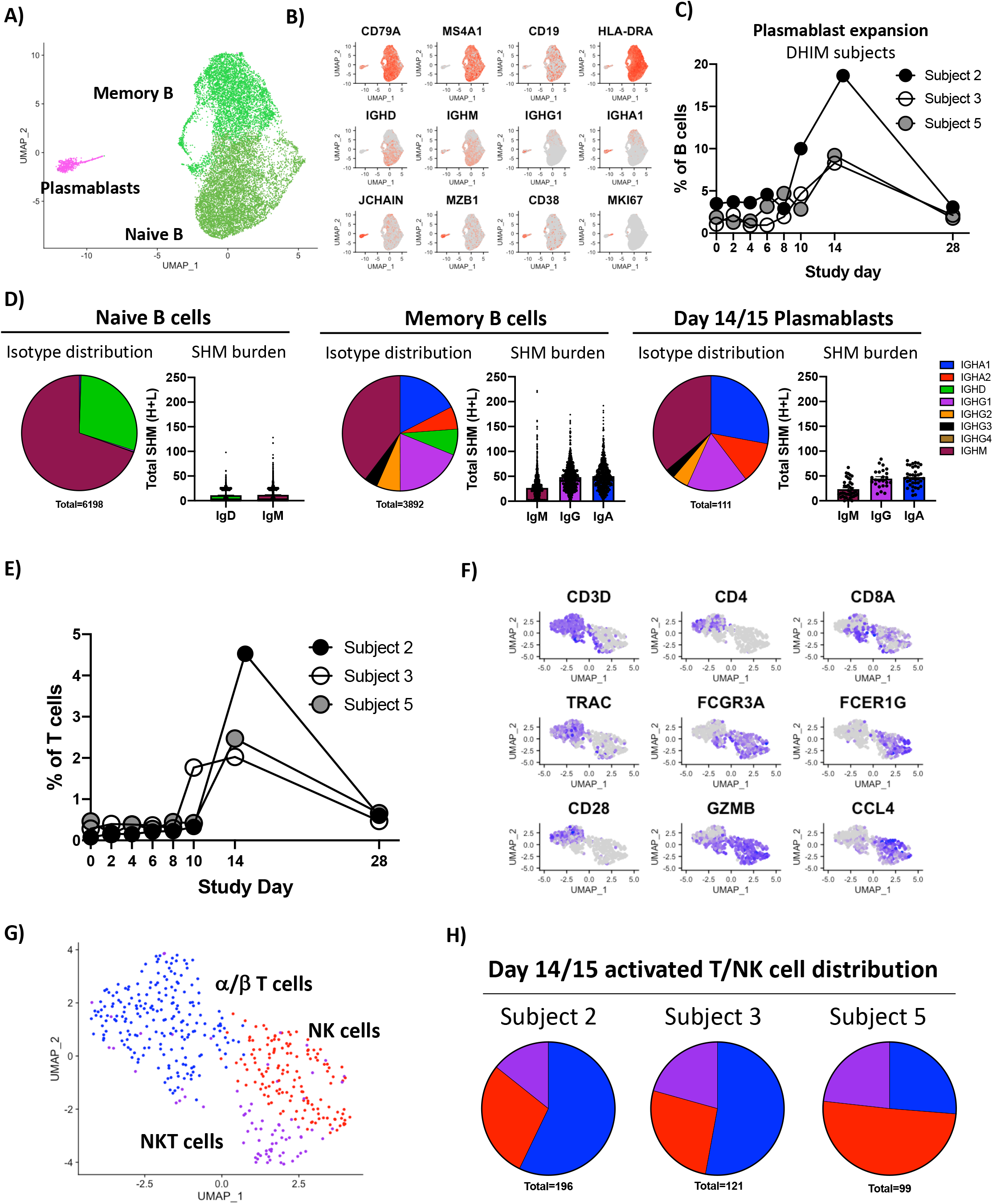
DHIM-associated lymphocyte expansion, quantification, and clonal characterization. **A)** Integrated UMAP projection of transcriptionally defined naïve B cells, memory B cells, and plasmablasts captured from all time points in all DHIM subjects. **B)** Expression of key linage-defining gene products at all time points from all DHIM subjects. **C)** Expansion of plasmablasts as a fraction of all B cells in response to DHIM challenge. **D)** Isotype distribution and hypermutation burden of transcriptionally defined naive B cells (all time points), memory B cells (all time points), and plasmablasts (study days 14/15). Analysis restricted to cells from which a single paired heavy/light chain was identified. **E)** Expansion of transcriptionally defined activated T/NK cells as a fraction of all T/NK cells in response to DHIM challenge. **F)** Expression of key activated T/NK cell gene products at all time points from all DHIM subjects at study day 14/15. **G)** Integrated UMAP projection of transcriptionally defined activated α/β T cells, activated NK cells, and activated NK/T cells from all DHIM subjects at study day 14/15. **I)** Distribution of transcriptionally defined activated α/β T cells, activated NK cells, and activated NK/T cells across all DHIM subjects at study day 14/15.

In addition to the plasmablast-phenotype B cells, a distinct transcriptional cluster of phenotypically activated T/NK cells was noted in all DHIM subjects in the scRNAseq analysis. This population expanded dramatically in relative frequency on DHIM study days 14/15 (**Figure 3E**), corresponding with the onset of T/NK cell activation as assessed by flow cytometry (**Supplemental Figure 7, Supplemental Figure 8**). To better understand the cell types involved in the cellular response to DHIM challenge, this population of activated T/NK cells was extracted into a new dataset and re-analyzed to allow for more granular assessment of cellular identities. Re-clustering of the cell contained within the activated T/NK cell population on DHIM study days 14/15 revealed 3 distinct transcriptional clusters (**Figure 3F, Figure 3G).** Expression of canonical T/NK cell gene products revealed that these clusters correspond to activated α/β T cells, activated NK cells, and activated NKT cells (**Figure 3F, Figure 3G**). These three activated T/NK populations were observed in all three DHIM subjects and mirrored the relative distribution of activated cells observed by flow cytometry (**Figure 3H, Supplemental Figure 7, Supplemental Figure 8**). These data suggest that DHIM challenge is associated with activation of both conventional α/β T cells, NK, cells, and NKT cells, albeit at reduced frequencies relative to what has been reported following natural primary DENV-1 infection (*13*).

### Identification of conserved DHIM gene signatures

To determine the kinetics and composition of the subject/population-specific transcriptional response to experimental DENV-1 infection, we performed differentially expressed gene (DEG) analysis across all DHIM samples. For this analysis, each annotated cell population on post-challenge study days was compared to the same population from the subject’s baseline sample (study day 0) using a Wilcoxon Log Rank test with a Bonferroni correction for multiple comparisons. Only genes with corrected p value of < 0.05 also exhibiting a >0.5 fold change in expression were considered significant. Although some modest and inconsistent transcriptional changes were observed on DHIM study day 6 and 8, it was at study day 10 that a consistent and robust transcriptional response to experimental DENV-1 infection was observed across all subjects (**Figure 4A**). The cellular populations exhibiting the most dramatic response to experimental DENV-1 infection were conventional and CD16^hi^ monocytes, with both populations containing > 100 DEGs on study day 10.

**Figure 4.**
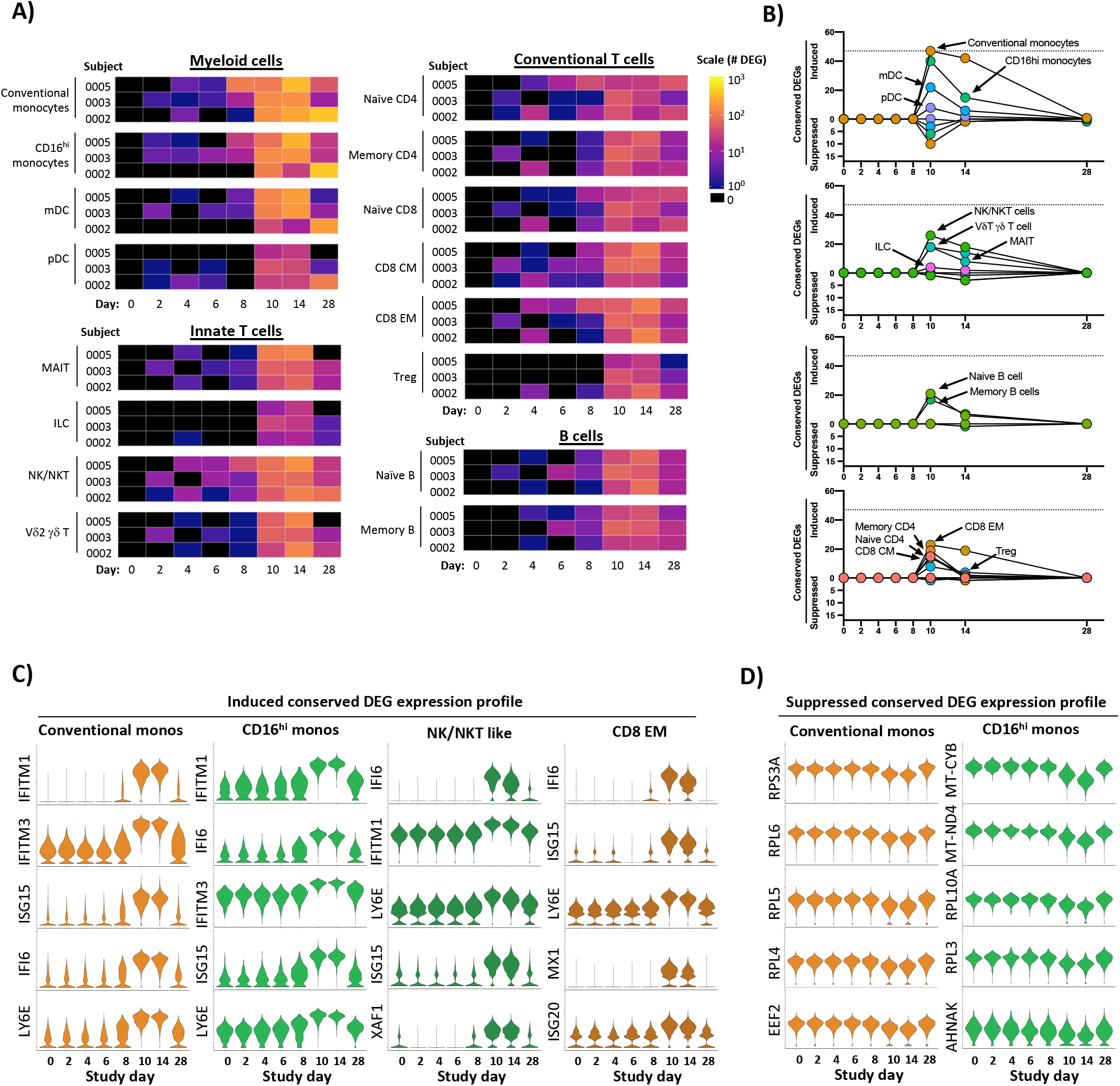
Temporal and transcriptional characterization of controlled DENV infection and identification of conserved gene signatures. **A)** Quantification of differential gene expression across all major leukocyte populations following experimental DENV infection. Subject- and population-specific differentially expressed genes (DEGs) were defined by a Wilcoxon rank-sum test with a Bonferroni correction relative to DHIM study day 0 for each subject. **B)** Population-restricted frequency and temporal dynamics of cDEGs across all DHIM subjects. Conserved DEGs defined as DEGs observed in all three subjects at the same time point relative to baseline. **C)** Expression of select upregulated cDEGs from DHIM study day 10 within the indicated cell populations across all study time points. **D)** Expression of select suppressed cDEGs from DHIM study day 10 within the indicated cell populations across all study time points.

To better define the conserved “core” transcriptional signature associated with experimental DENV-1 infection, we reduced our differential transcriptional analysis to only contain genes that were differently expressed from baseline in all three DHIM subjects at the indicated time point (**Figure 4B, Supplemental Table 7**). These conserved DEGs (cDEGs) were additionally separated into those induced by infection, and those suppressed by infection. This modified analysis again demonstrated that both conventional and CD16^hi^ monocytes exhibited the most robust conserved transcriptional response to experimental DENV-1 infection, followed in magnitude by NK/NKT cells, and CD8 Effector Memory (EM/) cells (**Figure 4B, Supplemental Table 7**). The majority of the annotated cDEGs were upregulated by DENV-1 infection, with only a minor population of cDEGs consistently suppressed in response to experimental DENV-1 infection (**Figure 4B, Supplemental Table 7**). The genes that were most consistently and significantly upregulated following experimental DENV-1 infection primarily corresponded to interferon-induced gene products (e.g. IFITM1, IFI6, ISG15, and MX1) and other genes associated with acute inflammation (e.g. TRIM22 and LY6E) (**Supplemental Table 7**). The expression of these gene products consistently peaked on study days 10 – 14, and trended towards baseline by study day 28 (**Figure 4C, Supplemental Table 7**). The few cDEGs that were repressed in response to experimental DENV-1 infection primarily corresponded to ribosomal protein subunits (e.g. RPL4, RPL5 and RPL6) translation elongation products (e.g. EEF2, EIF3L and EIF4B) mitochondrial associated gene products (e.g. MT-CYB and MT-ND4) (**Figure 4D, Supplemental Table 7**). This transcriptional profile is consistent with canonical IFN-associated suppression of protein translation and mitochondrial biogenesis and highlights a key mechanistic response to acute viral infection (*18, 19*). These conserved transcriptional responses to experimental DENV-1 infection not only reflect the acute cytokine-driven response to inflammation and infection but also capture the physiological response to inflammation and the induction of a distributed classic anti-viral cellular state in classical and CD16^hi^ monocytes.

### Identification of conserved natural primary DENV infection gene signatures

Having defined the core conserved transcriptional profile associated with experimental DENV-1 infection, we performed the same analysis on the cells from individuals experiencing a natural primary DENV-1 infection. For this analysis, each annotated cell population from a given subject’s acute infection samples (acute #1, acute #2) was compared to the same population from the subject’s 6-month reference sample (**Figure 5A**). As was observed in the DHIM subjects, the cellular populations exhibiting the most dramatic response to natural primary DENV-1 infection were conventional and CD16^hi^ monocytes, with both populations containing > 300 DEGs in both acute time points in both subjects. However, the number of population-specific DEGs observed in response to natural primary DENV-1 infection was several fold higher than the response observed in the same population following experimental DENV-1 infection (**Supplemental Figure 9**).

**Figure 5.**
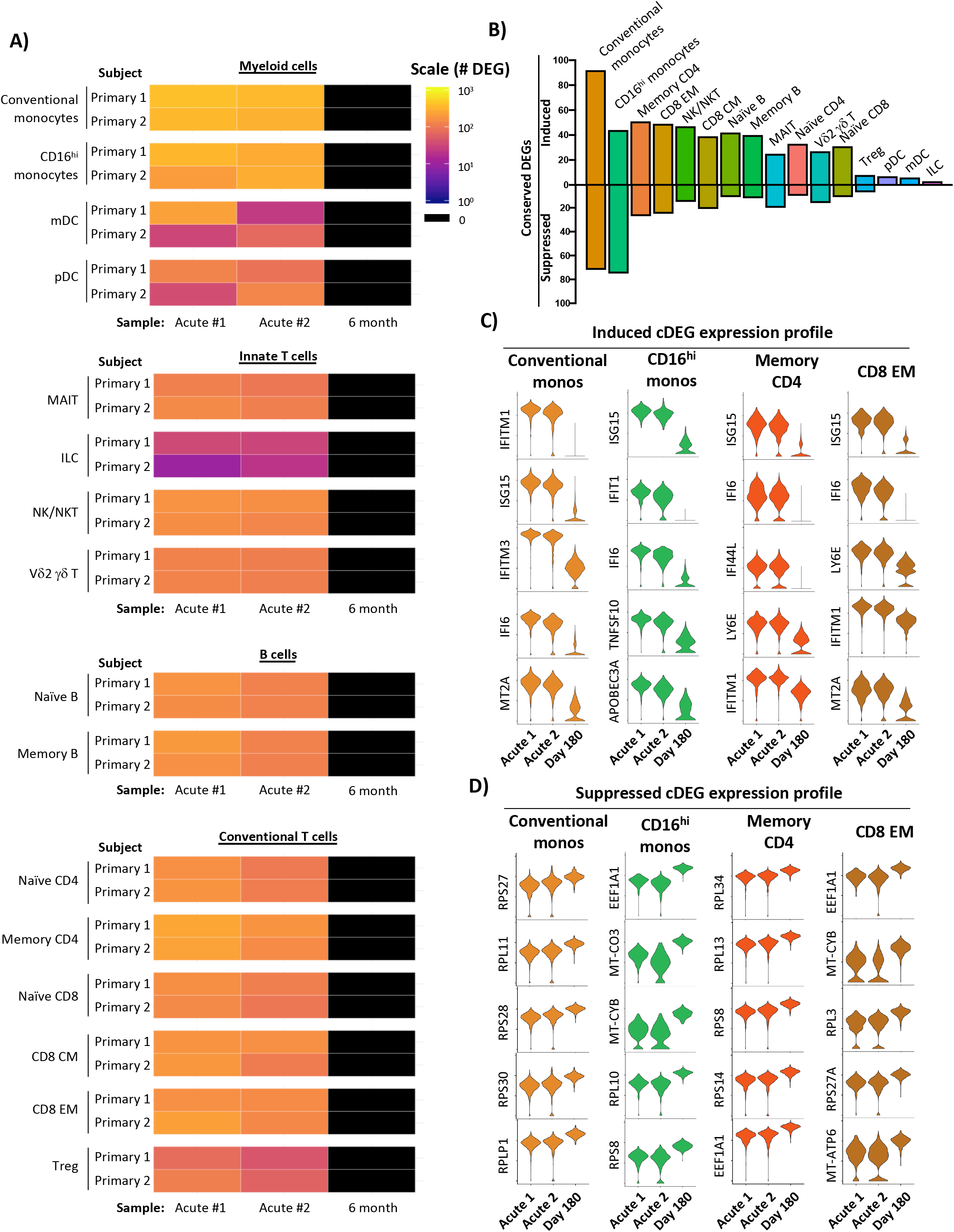
Temporal and transcriptional characterization of natural primary DENV infection and identification of conserved gene signatures. **A)** Quantification of differential gene expression across all major leukocyte populations following natural primary DENV-1 infection. Subject- and population-specific differentially expressed genes (DEGs) were defined by a Wilcoxon rank-sum test with a Bonferroni correction relative to the 6-month control sample for each subject. **B)** Population-restricted frequency of cDEGs from both natural primary DENV-1 infection subjects. **C)** Expression of select induced cDEGs within the indicated cell populations across all study time points. **D)** Expression of select induced cDEGs within the indicated cell populations across all study time points.

To reduce the complexity of the data, we again endeavored to define a core set of population-specific cDEGs within the natural primary DENV-1 infection dataset. As the samples obtained from natural primary DENV-1 infection were collected on consecutive days and the exact timing post-infection is unknown, cDEGs for these samples were defined as those gene products differentially expressed in all acute infection time points relative to their respective reference sample (the intersection of the acute timepoint DEGs). Conventional monocytes, memory CD4 T cells, and CD8 EM cells exhibited the largest number of upregulated cDEG across all annotated cell populations, with the majority of the conserved upregulated DEGs corresponding to classic interferon-induced gene products (e.g. IFIT1, IFI44L, and ISG20) and other genes associated with acute inflammation (e.g. TRIM22 and LY6E) (**Figure 5B, Figure 5C, Supplemental Table 8**). In contrast to the modest number of cDEGs suppressed in response to experimental DENV1 infection, a significant fraction of the cDEGs identified in natural primary DENV-1 infection were suppressed relative to baseline (**Figure 5B**). However, similarly to those suppressed conserved DEGs identified following experimental DENV-1 infection, the suppressed cDEGs identified following natural primary DENV-1 infection primarily corresponded to ribosomal protein subunits (e.g. RPL4, RPL5, and RPL6), translation elongation products (e.g. EEF2, EIF3L, and EIF4B) and mitochondrial associated gene products (e.g. MT-CYB, and MT-ND4) (**Figure 5D, Supplemental Table 8)**. These data suggest that natural primary DENV infection not only induces the expression of interferon-induced gene products in a wide range of cell type, but that the physiological consequence this response is profound and widely distributed.

#### Transcriptional profile overlap between controlled and natural primary DENV infection

Having identified cell-specific conserved sets of differently expressed genes present in PBMC subpopulations following experimental and natural primary DENV infection, we assessed the overlap between these two datasets to compare the conserved transcriptional response to experimental and natural primary DENV infection. In light of the fact that cells obtained 10 days post infection in the DHIM study contained the most cDEGs relative to baseline, this time point was selected for comparison against the natural primary infection dataset.

Although the overall number of infection-induced cDEGs identified in cells from natural primary DENV-1 infection was greater than that observed following experimental DENV-1 infection, the cell population specific frequencies of infection-induced cDEGs in each population were highly correlated between groups (R^2^ = 0.6825, p < 0.0001) (**Figure 6A**). Furthermore, the infection-induced cDEGs identified following experimental DENV were determined to consistently represent a subset of the cDEGs induced in natural primary DENV-1 infection (**Figure 6B, Table 1**). Gene ontogeny analysis revealed that the infection-induced cDEGs that were found in common to experimental and natural primary DENV infection datasets fell into classic interferon response pathways and negative regulation of viral replication (**Table 2**). Those cDEGs which were unique to natural primary DENV-1 infection corresponded to gene pathways involved in cytokine secretion, antigen processing/presentation, and detection of viruses (**Table 2**).

**Table 1.** Conserved and unique infection-induced cDEG identified following natural infection or on DHIM study day 10

| Population | Unique DHIM | Common | Unique Natural infection |
| --- | --- | --- | --- |
| **Conventional monocytes** | HBB, HES4, NCF1, STAT2, TNFSF13B, TXNIP, WARS | APOBEC3A, EIF2AK2, EPSTI1, GBP1, IFI35, IFI44, IFI44L, IFI6, IFITM1, IFITM2, IFITM3, IRF7, ISG15, ISG20, LAP3, LY6E, MT2A, MX1, MX2, OAS1, OAS3, OASL, PARP14, PLAC8, PLSCR1, PSMB9, PSME2, RNF213, RSAD2, SAMD4A, SERPING1, SIGLEC1, STAT1, TMEM123, TNFSF10, TRIM22, TYMP, UBE2L6, VAMP5, XAF1 | BST2, CCL2, CCR1, CMPK2, CTSL, CXCL10, DDX58, DDX60L, DRAP1, FCGR1A, GBP4, GBP5, GCH1, GIMAP4, HERC5, HLA-A, HLA-B, HLA-C, HSH2D, IFI16, IFI27, IFIH1, IFIT1, IFIT2, IFIT3, IL1RN, LGALS3BP, LGALS9, MAFB, MARCKS, MYL12A, NAPA, NMI, NT5C3A, OAS2, PARP12, PARP9, PHF11, PML, PSMA4, RNASE2, SAMD9, SAMD9L, SCO2, SELL, SMCHD1, SP110, SPATS2L, TCN2, USP18, XRN1, ZBP1 |
| **CD16hi monocytes** | BST2, GBP1, HES4, IFITM1, IFITM3, LAP3, LY6E, OAS1, PARP14, PLAC8, PSMB9, PSME2, TMEM123, TNFSF13B, TYMP, UBE2L6, VAMP5, WARS | APOBEC3A, CXCL10, EIF2AK2, EPSTI1, HERC5, IFI35, IFI44, IFI44L, IFI6, IFIT2, IRF7, ISG15, MX1, MX2, NCF1, OAS3, PLSCR1, RSAD2, SERPING1, TNFSF10, TRIM22, XAF1 | CD300E, CDKN1A, DDX58, DDX60L, FFAR2, GBP4, GLUL, HLA-A, IFI27, IFIH1, IFIT1, IFIT3, LGALS3BP, LGALS9, MARCKS, OASL, SAMD9L, SIGLEC1, SPATS2L, TCN2, USP18, ZBP1 |
| **CD4 memory** | IFITM2 | BST2, EIF2AK2, EPSTI1, IFI44, IFI44L, IFI6, IFITM1, ISG15, ISG20, LY6E, MX2, PARP9, PLSCR1, PSMB9, SP100, STAT1, TRIM22, XAF1 | ADAR, DRAP1, GBP1, HERC5, HLA-A, HLA-B, HLA-E, IER2, IFI16, IFI35, IFIT3, IFITM3, IRF7, LGALS9, MT2A, MX1, OAS1, OAS3, OASL, PARP10, PARP12, PHF11, PSME2, RNF213, RSAD2, SAMD9, SAMD9L, SMCHD1, SP110, TNFSF10, TYMP, UBE2L6, ZBP1 |
| **NK/NKT** | IFITM2, LGALS1, PSMB9, PSME2, SHISA5 | BST2, EIF2AK2, EPSTI1, IFI35, IFI44L, IFI6, IFITM1, IFITM3, IRF7, ISG15, ISG20, LY6E, MX1, MX2, PARP9, PLSCR1, STAT1, TRIM22, TYMP, UBE2L6, XAF1 | ADAR, CD38, DTX3L, GBP1, IFI16, IFI44, IFIT1, IFIT3, LAG3, LGALS9, MT2A, NT5C3A, OAS1, OAS2, OAS3, OASL, PARP12, PRF1, RNF213, RSAD2, S100A11, SAMD9, SAMD9L, SP100, WARS, ZBP1 |
| **CD8 EM** | IFITM2 | BST2, DRAP1, EIF2AK2, IFI35, IFI44L, IFI6, IFITM1, IFITM3, IRF7, ISG15, ISG20, LY6E, MT2A, MX1, MX2, PLSCR1, PSMB9, PSME2, SP100, STAT1, TRIM22, XAF1 | ADAR, CD38, EPSTI1, GBP1, HERC5, IFI16, IFI44, IFIT1, IFIT3, LAG3, LGALS9, OAS1, OAS2, OASL, PARP10, PARP9, PRF1, RNF213, RSAD2, S100A11, SAMD9, SAMD9L, SP110, TAP1, TYMP, UBE2L6 |

**Table 2.** Enriched GO terms in overlapping and natural infection only cell populations

| DHIM AND Natural Infection |  |  |  |  |  | Natural Infection ONLY |  |  |  |
| --- | --- | --- | --- | --- | --- | --- | --- | --- | --- |
| Cell type | Rank | GO term | Description | FE | FDR | GO term | Description | FE | FDR |
| Conventional mono | 1 | 0032020 | ISG15-protein conjugation | > 100 | 3.20E-02 | 0034344 | regulation of type III interferon production | > 100 | 1.30E-02 |
| Conventional mono | 2 | 0060700 | regulation of ribonuclease activity | > 100 | 5.92E-04 | 0035549 | positive regulation of interferon-beta secretion | > 100 | 2.42E-02 |
| Conventional mono | 3 | 0016185 | synaptic vesicle budding from presynaptic endocytic zone membrane | > 100 | 3.95E-02 | 0009597 | detection of virus | > 100 | 3.06E-02 |
| Conventional mono | 4 | 0035455 | response to interferon-alpha | > 100 | 1.15E-08 | 0035547 | regulation of interferon-beta secretion | > 100 | 3.03E-02 |
| Conventional mono | 5 | 0060337 | type I interferon signaling pathway | > 100 | 6.28E-25 | 0039528 | cytoplasmic pattern recognition receptor signaling pathway in response to virus | > 100 | 3.00E-02 |
| Conventional mono | 6 | 0071357 | cellular response to type I interferon | > 100 | 4.19E-25 | 0002480 | antigen processing and presentation of exogenous peptide antigen via MHC class I, TAP-independent | > 100 | 1.29E-03 |
| Conventional mono | 7 | 0034340 | response to type I interferon | > 100 | 8.74E-25 | 0002486 | antigen processing and presentation of endogenous peptide antigen via MHC class I via ER pathway, TAP-independent | 94.78 | 2.35E-03 |
| Conventional mono | 8 | 0045071 | negative regulation of viral genome replication | > 100 | 2.03E-19 | 0002484 | antigen processing and presentation of endogenous peptide antigen via MHC class I via ER pathway | 94.78 | 2.30E-03 |
| CD16<sup>hi</sup> monos | 1 | 0032020 | ISG15-protein conjugation | > 100 | 1.27E-02 | 0034344 | regulation of type III interferon production | > 100 | 6.83E-03 |
| CD16<sup>hi</sup> monos | 2 | 0016185 | synaptic vesicle budding from presynaptic endocytic zone membrane | > 100 | 1.59E-02 | 0035549 | positive regulation of interferon-beta secretion | > 100 | 1.11E-02 |
| CD16<sup>hi</sup> monos | 3 | 0140239 | postsynaptic endocytosis | > 100 | 2.31E-02 | 0009597 | detection of virus | > 100 | 1.29E-02 |
| CD16<sup>hi</sup> monos | 4 | 0098884 | postsynaptic neurotransmitter receptor internalization | > 100 | 2.25E-02 | 0035547 | regulation of interferon-beta secretion | > 100 | 1.26E-02 |
| CD16<sup>hi</sup> monos | 5 | 0070142 | synaptic vesicle budding | > 100 | 2.46E-02 | 0039528 | cytoplasmic pattern recognition receptor signaling pathway in response to virus | > 100 | 1.23E-02 |
| CD16<sup>hi</sup> monos | 6 | 0060337 | type I interferon signaling pathway | > 100 | 3.16E-16 | 1902741 | positive regulation of interferon-alpha secretion | > 100 | 2.56E-02 |
| CD16<sup>hi</sup> monos | 7 | 0071357 | cellular response to type I interferon | > 100 | 2.81E-16 | 1902739 | regulation of interferon-alpha secretion | > 100 | 2.51E-02 |
| CD16<sup>hi</sup> monos | 8 | 0099590 | neurotransmitter receptor internalization | > 100 | 4.18E-02 | 0071360 | cellular response to exogenous dsRNA | > 100 | 1.53E-03 |
| Memory CD4 | 1 | 0035455 | response to interferon-alpha | > 100 | 4.30E-06 | 0002519 | natural killer cell tolerance induction | > 100 | 5.10E-03 |
| Memory CD4 | 2 | 0035455 | response to interferon-beta | > 100 | 7.09E-08 | 0042270 | protection from natural killer cell mediated cytotoxicity | > 100 | 9.10E-03 |
| Memory CD4 | 3 | 0060337 | type I interferon signaling pathway | > 100 | 1.73E-14 | 0002480 | antigen processing and presentation of exogenous peptide antigen via MHC class I, TAP-independent | > 100 | 3.11E-04 |
| Memory CD4 | 4 | 0071357 | cellular response to type I interferon | > 100 | 1.30E-14 | 0060700 | regulation of ribonuclease activity | > 100 | 3.04E-04 |
| Memory CD4 | 5 | 0034340 | response to type I interferon | > 100 | 1.90E-14 | 0032020 | ISG15-protein conjugation | > 100 | 1.17E-02 |
| Memory CD4 | 6 | 0045071 | negative regulation of viral genome replication | > 100 | 1.24E-08 | 0002486 | antigen processing and presentation of endogenous peptide antigen via MHC class I via ER pathway, TAP-independent | > 100 | 5.99E-04 |
| Memory CD4 | 7 | 0048525 | negative regulation of viral process | 89.11 | 3.23E-11 | 0002484 | antigen processing and presentation of endogenous peptide antigen via MHC class I via ER pathway | > 100 | 5.87E-04 |
| Memory CD4 | 8 | 1903901 | negative regulation of viral life cycle | 81.77 | 7.16E-08 | 2001187 | positive regulation of CD8-positive, alpha-beta T cell activation | > 100 | 1.71E-02 |
| NK/NKT | 1 | 0016185 | synaptic vesicle budding from presynaptic endocytic zone membrane | > 100 | 1.02E-02 | 0060700 | regulation of ribonuclease activity | > 100 | 8.34E-07 |
| NK/NKT | 2 | 0140239 | postsynaptic endocytosis | > 100 | 1.28E-02 | 0032069 | regulation of nuclease activity | > 100 | 1.62E-05 |
| NK/NKT | 3 | 0098884 | postsynaptic neurotransmitter receptor internalization | > 100 | 1.91E-02 | 0060337 | type I interferon signaling pathway | > 100 | 1.22E-12 |
| NK/NKT | 4 | 0035455 | response to interferon-alpha | > 100 | 1.87E-02 | 0071357 | cellular response to type I interferon | > 100 | 9.73E-13 |
| NK/NKT | 5 | 0070142 | synaptic vesicle budding | > 100 | 3.96E-08 | 0035455 | response to interferon-alpha | 96.53 | 2.38E-03 |
| NK/NKT | 6 | 0035456 | response to interferon-beta | > 100 | 2.14E-02 | 0034340 | response to type I interferon | 96.53 | 1.48E-12 |
| NK/NKT | 7 | 0060337 | type I interferon signaling pathway | > 100 | 8.49E-10 | 0045071 | negative regulation of viral genome replication | 87.19 | 2.11E-09 |
| NK/NKT | 8 | 0071357 | cellular response to type I interferon | > 100 | 8.20E-21 | 0060333 | interferon-gamma-mediated signaling pathway | 75.08 | 5.36E-09 |
| CD8 EM | 1 | 0016185 | synaptic vesicle budding from presynaptic endocytic zone membrane | > 100 | 1.21E-02 | 0060700 | regulation of ribonuclease activity | > 100 | 1.65E-04 |
| CD8 EM | 2 | 0140239 | postsynaptic endocytosis | > 100 | 1.77E-02 | 0016185 | synaptic vesicle budding from presynaptic endocytic zone membrane | > 100 | 1.54E-02 |
| CD8 EM | 3 | 0098884 | postsynaptic neurotransmitter receptor internalization | > 100 | 1.74E-02 | 0140239 | postsynaptic endocytosis | > 100 | 2.26E-02 |
| CD8 EM | 4 | 0035455 | response to interferon-alpha | > 100 | 4.36E-08 | 0098884 | postsynaptic neurotransmitter receptor internalization | > 100 | 2.22E-02 |
| CD8 EM | 5 | 0070142 | synaptic vesicle budding | > 100 | 1.94E-02 | 0070142 | synaptic vesicle budding | > 100 | 2.61E-02 |
| CD8 EM | 6 | 0060337 | type I interferon signaling pathway | > 100 | 1.06E-22 | 0035455 | response to interferon-alpha | > 100 | 1.76E-05 |
| CD8 EM | 7 | 0071357 | cellular response to type I interferon | > 100 | 5.32E-23 | 0060340 | positive regulation of type I interferon-mediated signaling pathway | > 100 | 3.93E-02 |
| CD8 EM | 8 | 0035456 | response to interferon-beta | > 100 | 9.32E-10 | 0035457 | cellular response to interferon-alpha | > 100 | 3.86E-02 |

**Figure 6.**
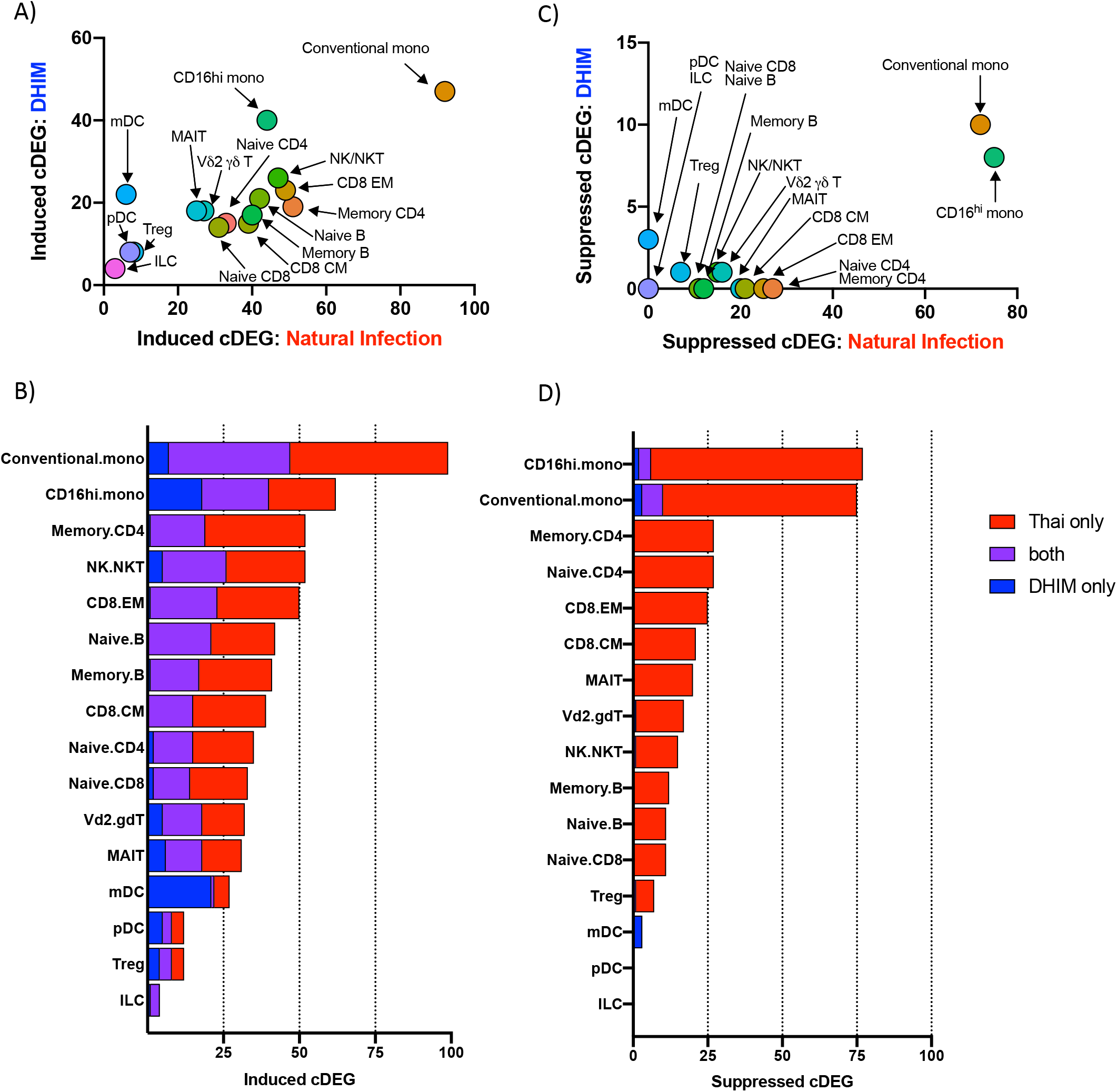
Transcriptional profile overlap of controlled and natural primary DENV-1 infection. **A)** Frequency and cell-population distribution of infection-induced cDEGs identified following natural primary or experimental DENV-1 infection. DHIM cDEGs restricted to study day 10. **B)** Frequency and overlap of infection-induced cDEGs identified following natural primary or experimental DENV-1 infection **C)** Frequency and cell-population distribution of infection-suppressed cDEGs identified following natural primary or experimental DENV-1 infection. DHIM cDEGs restricted to study day 10. **D)** Frequency and overlap of infection-suppressed cDEGs identified following natural primary or experimental DENV-1 infection

In contrast to the high degree of correlation observed in the cell-population specific frequencies of infection-induced cDEGs, infection-suppressed cDEGs were overwhelming restricted to those cells obtained following natural primary DENV infection (**Figure 6C**). However, the few infection-suppressed cDEGs observed in cells following experimental DENV infection generally represented a subset of those suppressed following natural primary DENV infection (**Figure 6D, Table 3**). Infection-suppressed cDEGs overwhelming represented gene products associated with protein translation/elongation, as well as mitochondrial function and biogenesis (**Table 3, Supplemental Table 8**). These data suggest that both natural and experimental primary DENV-1 infection upregulate a similar acute transcriptional program in similar cells, but that the greater magnitude of the that response elicited by natural primary DENV-1 infections has a significantly more profound physiological response on basic cellular processes, broadly suppressing protein translation and mitochondrial function. Furthermore, these data indicate that monocytes are uniquely responsive to both experimental and natural primary DENV infection.

**Table 3.** Conserved and unique infection-suppressed cDEG identified following natural infection or on DHIM study day 10

| Population | Unique DHIM | Common | Unique Natural infection |
| --- | --- | --- | --- |
| CD16hi monocytes | IL1B, AHNAK | RPL10A, RPL3, MT-ND4, MT-CYB | RPL6, RPS3A, EEF1A1, RPS15A, RPL10, RPS13, RPS8, RPS23, RPL34, RPS4X, RPS27A, RPL8, RPL14, RPS14, RPS7, RPL18A, RPS6, RPL12, RPLP2, RPL35A, RPL18, RPL11, RPL21, RPL7A, RPL9, RPL5, SLC25A6, RPL32, RPS24, RPS21, RPL13, RPL30, RPL4, MT-CO3, RPL19, RPL22, RPS16, RPL26, RPL15, GNB2L1, COTL1, PABPC1, SLC25A5, ACTG1, EEF2, EEF1B2, NACA, RPLP0, RPL27, RPS25, RPS5, TPT1, RPL23A, RPS3, RPSA, RPS18, PFDN5, EIF3K, RPL37A, UQCRB, BTF3, RPL29, MT-ND3, RPS4Y1, RPL23, YBX1, CCNI, RPS17, NAP1L1, EIF3L, RBM3 |
| Conventional monocytes | IL1B, CXCL8, EIF4B | RPS3A, RPL4, EEF2, RPL6, RPL5, EIF3L, RPL3 | TPT1, RPS28, EEF1A1, RPS13, RPLP1, RPL8, RPL18, RPS7, RPS14, RPL34, RPS24, RPL35A, RPS15A, MT-ND3, MT-CO3, RPS8, RPS4X, RPL11, RPL10, RPL19, RPL9, NACA, RPL22, RPL26, RPL37, RPS23, RPL15, RPL7A, RPL32, RPL18A, BTF3, RPL30, RPLP2, RPS16, RPS27A, RPS6, RPLP0, RPL29, COX4I1, PABPC1, CSTA, COTL1, SLC25A6, RPL27, RPL13, RPL10A, RPS5, EEF1B2, RPL21, AP1S2, RPS3, GSTP1, RPS18, PCBP2, RPL7, GNB2L1, SLC25A5, RPS4Y1, ALDH2, EIF3F, RBM3, FCGRT, EIF3E, EIF3H, CRTAP |

## DISCUSSION

In this study, we utilized high-throughput single-cell RNA sequencing (scRNAseq) technology to assess the longitudinal transcriptional profile in both controlled and natural primary DENV-1 infection with single-cell resolution. Core sets of conserved Differentially Express Genes that were induced or suppressed by either natural or experimental primary DENV were identified, and the overlap between the study groups assessed. The infection-induced cDEGs associated with experimental DENV infection were found to reflect a subset within the larger gene set associated with natural primary DENV infection, primarily corresponding to gene products associated with a cellular response to systemic inflammation and interferon production. In contrast, infection-suppressed cDEGs were much more common in cells obtained from natural primary DENV infection than in cells obtained from experimental DENV infection. Infection-suppressed cDEGs primarily corresponded to gene products associated with protein translation/elongation and mitochondrial function, two cellular processes known to be suppressed by IFN signaling. These results are consistent with the concept that the immune response elicited by DHIM represents a tempered version of that generated in response to a natural primary DENV infection, but that the more pronounced inflammation associated with natural primary DENV infection has a correspondingly more pronounced impact on basic cellular processes to induce a multi-layered systemic anti-viral state. These data provide insight into the molecular level response to DENV infection, and how viral pathogenesis correlates with immune activation and cellular pathophysiology.

Interferon-induced gene products render cells inhospitable to viral replication and propagation through multiple mechanisms. These includes the expression of gene products that directly degrade viral RNA/DNA, the post-translational inhibition of viral entry and/or budding, post-translation inhibition of protein translation, as well as restriction of key metabolites and macromolecules required for virus replication and virion assembly (*20*). Transcriptionally repressing the expression of gene products necessary for protein translation/elongation and mitochondrial function is one of the less specific – though highly effective – methods by which IFN exposure can limit viral replication (*18, 20*). This process of broadly inhibiting a function so central to cellular survival is understandably tightly regulated, and subject to multiple homeostatic feedback loops. However, the data presented here suggests that a tipping point may exist, where the magnitude of the interferon-induced transcriptional response reaches a threshold, whereupon a broader range of cellular factors are significantly impacted causing a qualitative change in transcriptional response to infection.

The observation that both the induction of IFN-responsive gene products as well as the suppression of genes associated with protein translation and mitochondrial function was most dramatically and consistently observed in conventional/CD16^hi^ monocytes may help explain certain discrepancies in the literature regarding DENV cellular tropism. Historically, monocytes, macrophages, and dendritic cells have been cited as the primary circulating cellular reservoir of DENV (*21, 22*). However, this claim was primarily based on *in vitro* infection experiments, where the impact of systemic inflammation and interferon signaling may not impact cellular permissiveness to infection. More recent studies utilizing flow cytometry, RT-PCR, and scRNAseq analysis have suggested that B cells are the primary circulating natural primary reservoir of DENV (*23–28*). Our observation that monocytes appear to much more dramatically respond to DENV-elicited systemic inflammation than B cells may suggest that while monocytes are more permissive to infection *in vitro*, they become a much less attractive target of infection and replication *in vivo*.

Although the data presented herein represents the most intensive and high-resolution transcriptional assessment of both natural and experimental primary DENV infection to date, several limitations of the study must be acknowledged. Firstly, the relatively small numbers of subjects analyzed in both arms of the study and the significant age and background differences between the subjects are potential confounders for the analysis. In addition to viral genetic differences, the route and method of virus inoculation differs significantly between natural and experimental primary DENV infection (mosquito vectored vs intradermal needle). The impact of these variables on the immunological profile described here can only be addressed with future studies.

In their totality, the data presented in this study suggest that the immunological profile elicited by the administration of an attenuated DENV in a controlled setting activates many of the same molecular pathways in the same cells as unattenuated, naturally-acquired primary DENV. This includes the activation and expansion of lymphocyte populations (T/NK/NKT cells and plasmablasts), as well as predictable perturbation in the transcriptional profile of all major leukocyte subsets. Furthermore, it highlights the unique role that monocytes play in responding to acute DENV infection, potentially offering a framework to develop more accurate and rapid diagnostic metrics in the future.

## MATERIALS AND METHODS

### Dengue Human Challenge Model

Peripheral blood mononuclear cells (PBMC) and plasma for this study were obtained from the previously described phase 1 open label, Dengue Virus-1 Live Virus Human Challenge (DENV-1-LVHC) study performed at the State University of New York, Upstate Medical University in Syracuse, NY (*10*). The study was approved by the State University of New York Upstate Medical University (SUNY-UMU) and the Department of Defense’s Human Right Protection Organization (HRPO). ClinicalTrials.gov identifier for this trial is NCT02372175. All subjects included in this study received 3.25 × 10^3^ PFU of the 45AZ5 DENV-1 challenge strain virus following pre-screening to ensure an absence of preexisting DENV immunity. Flavivirus antibody screening, dengue IgM and IgG ELISA, microneutralization assay and quantitative reverse-transcriptase polymerase chain reaction (RT-PCR) were performed at the Viral Diseases Branch, Walter Reed Army Institute of Research (WRAIR) in Silver Spring, Maryland using previously published techniques(*29–32*).

### Natural primary DENV infection sample collection

PBMC and plasma were isolated from whole blood specimens obtained from children enrolled in a hospital-based acute febrile illness study in Bangkok, Thailand, the design of which has been previously described (*33, 34*). In brief, the study enrolled children who presented to the hospital with acute febrile illness. Blood samples were obtained daily during illness and at early and late convalescent time points; the term ‘fever day’ is used to report acute illness time points relative to Day 0, defined as the day of defervescence. The infecting virus type (DENV-1-4) was determined by RT-PCR and/or virus isolation as previously described (*35*), and serology (EIA and HAI assays) was used to distinguish primary and secondary DENV infections (*36*). Written informed consent was obtained from each subject and/or his/her parent or guardian. The study protocol was approved by the Institutional Review Boards of the Thai Ministry of Public Health, the Office of the U.S. Army Surgeon General, and the University of Massachusetts Medical School. PBMC and plasma samples were cryopreserved for later analysis.

### Flow Cytometry

Cryopreserved PBMC were thawed and placed in RPMI 1640 medium supplemented with 10% heat-inactivated fetal bovine serum, L-glutamine, penicillin, and streptomycin prior to analysis. Cell viability was assessed using CTL-LDC Dye (Cellular Technology Limited [CTL], Shaker Heights, OH) and a CTL-ImmunoSpot S6 Ultimate-V Analyzer (CTL). Surface staining for flow cytometry analysis was performed in PBS supplemented with 2% FBS at room temperature. Aqua Live/Dead or Violet Live/Dead dye (ThermoFisher) was used to exclude dead cells in all experiments. Antibodies and dilutions used for flow cytometry analysis are listed in **Supplemental Table 9.** Flow cytometry analysis was performed on a custom-order BD LSRFortessa instrument, and data analyzed using FlowJo v10.2 software (Treestar).

### Single-cell RNA sequencing library generation

Thawed PBMC suspensions were prepared for single-cell RNA sequencing using the Chromium Single-Cell 5′ Reagent version 2 kit and Chromium Single-Cell Controller (10x Genomics, CA)(*37*). 2000–8000 cells per reaction suspended at a density of 50–500 cells/μL in PBS plus 0.5% FBS were loaded for gel bead-in-emulsion (GEM) generation and barcoding. Reverse transcription, RT-cleanup, and cDNA amplification were performed to isolate and amplify cDNA for downstream 5′ gene or enriched V(D)J library construction according to the manufacturer’s protocol. Libraries were constructed using the Chromium Single-Cell 5′ reagent kit, V(D)J Human B Cell Enrichment Kit, 3′/5′ Library Construction Kit, and i7 Multiplex Kit (10x Genomics, CA) according to the manufacturer’s protocol.

### Sequencing

scRNAseq 5′ gene expression libraries and BCR V(D)J enriched libraries were sequenced on an Illumina NovaSeq 6000 instrument using the S2, or S4 reagent kits (300 cycles). Libraries were balanced to allow for ~150,000 reads/cell for 5′ gene expression libraries, and ~20,000 reads/cell for BCR V(D)J enriched libraries. Sequencing parameters were set for 150 cycles for Read1, 8 cycles for Index1, and 150 cycles for Read2. Prior to sequencing, library quality and concentration were assessed using an Agilent 4200 TapeStation with High Sensitivity D5000 ScreenTape Assay and Qubit Fluorometer (Thermo Fisher Scientific) with dsDNA BR assay kit according to the manufacturer’s recommendations

### 5’ gene expression analysis/visualization

5′ gene expression alignment from all PBMC samples was performed using the 10x Genomics Cell Ranger pipeline(*37*). Sample demultiplexing, alignment, barcode/UMI filtering, and duplicate compression was performed using the Cell Ranger software package (10x Genomics, CA, v2.1.0) and bcl2fastq2 (Illumina, CA, v2.20) according to the manufacturer’s recommendations, using the default settings and mkfastq/count commands, respectively. Transcript alignment was performed against a human reference library generated using the Cell Ranger mkref command, the Ensembl GRCh38 v87 top-level genome FASTA, and the corresponding Ensembl v87 gene GTF.

Multi-sample integration, data normalization, dimensional reduction, visualization, and differential gene expression were performed using the R package Seurat (v3.1.4) (*38, 39*). All datasets were filtered to only contain cells with between 200-6,000 unique features and <10% mitochondrial RNA content. To eliminate erythrocyte contamination, datasets were additionally filtered to contain cells with less than a 5% erythrocytic gene signature (defined as HBA1, HBA2, HBB). Data were scaled, transformed, and variable genes were identified using the SCTransform() function. SelectIntegrationFeatures() and PrepSCTIntegration() functions were used to identify conserved features for dataset integration, and final dataset anchoring/integration were performed using FindIntegrationAnchors() and IntegrateData() functions, with the day 0 DHIM samples and day 180 natural primary infection samples used as reference datasets. PCA was performed using variable genes defined by SCTransform() additionally filtered to remove TCR V/D/J or BCR κ/λ gene segments. The first 33 resultant PCs were used to perform a UMAP dimensional reduction of the dataset (RunUMAP()) and to construct a shared nearest neighbor graph (SNN; FindNeighbors()). This SNN was used to cluster the dataset (FindClusters()) with default parameters and resolution set to 0.4.

Following dataset integration and dimensional reduction/clustering, gene expression data was log_e_(UMI+1) transformed and scaled by a factor of 10,000 using the NormalizeData() function. This normalized gene expression data was used to determine cellular cluster identity by utilizing the Seurat application of a Wilcoxon rank-sum test (FindAllMarkers()), and comparing the resulting differential expression data to known cell-linage specific gene sets. Differential gene expression analysis between study time points was performed using normalized gene expression data and the Wilcoxon rank-sum test with implementation in the FindMarkers() function, with a log_e_ fold chain threshold of 0.5 and min.pct of 0.25. Bonferroni correction was used to control for Fall Discovery Rate (FDR), with a corrected p value of < 0.05 considered significant.

### BCR sequence analysis

BCR clonotype identification, alignment, and annotation were performed using the 10x Genomics Cell Ranger pipeline. Sample demultiplexing and clonotype alignment was performed using the Cell Ranger software package (10x Genomics, CA, v2.1.0) and bcl2fastq2 (Illumina, CA, v2.20) according to the manufacturer’s recommendations, using the default settings and mkfastq/vdj commands, respectively. Immunoglobulin clonotype alignment was performed against a filtered human V(D)J reference library generated using the Cell Ranger mkvdjref command and the Ensembl GRCh38 v87 top-level genome FASTA and the corresponding Ensembl v87 gene GTF. BCR hypermutation burden was assessed using the software package BRILIA(*40*)

### Recombinant monoclonal antibody synthesis

The variable regions from the heavy and light chains of targeted immunoglobulin sequences were codon optimized, synthesized *in vitro* and subcloned into a pcDNA3.4 vector containing the human IgG1 Fc region by a commercial partner (Genscript). Transfection grade plasmids were purified by maxiprep and transfected into a 293-6E expression system. Cells were grown in serum-free FreeStyle 293 Expression Media (Thermo Fisher), and the cell supernatants collected on day 6 for antibody purification. Following centrifugation and filtration, the cell culture supernatant was loaded onto an affinity purification column, washed, eluted, and buffer exchanged to the final formulation buffer (PBS). Antibody lot purity was assessed by SDS-PAGE, and the final concentration determined by 280nm absorption. The clonotype information for all monoclonal antibodies generated as part of this study is listed in **Supplemental Table 5**.

### Viruses

DENV1-4 (strains Western Pacific 1974, S16803, CH53489, and TVP-360, respectively) propagated in C6/36 mosquito cells were utilized for ELISA. Virus for ELISA was purified by ultracentrifugation through a 30% sucrose solution and the virus pellet was resuspended in PBS.

### Monoclonal antibody DENV-capture ELISA

Monoclonal antibody DENV reactivity was assessed using a 4G2 DENV capture ELISA protocol. In short, 96 well NUNC MaxSorb flat-bottom plates were coated with 2 μg/ml flavivirus group-reactive mouse monoclonal antibody 4G2 (Envigo Bioproducts, Inc.) diluted in borate saline buffer. Plates were washed and blocked with 0.25% BSA + 1% Normal Goat Serum in PBS after overnight incubation. DENV-1, −2, −3 or −4 (strains Western Pacific 1974, S16803, CH53489, and TVP-360, respectively) were captured for 2 hours in the appropriate wells, followed by extensive washing. Serially diluted monoclonal antibody samples were incubated for 1hr at RT on the captured virus, and DENV-specific antibody binding quantified using anti-human IgG HRP (Sigma-Aldrich, SAB3701362). Secondary antibody binding was quantified using the TMB Microwell Peroxidase Substrate System (KPL, cat. #50-76-00) and Synergy HT plate reader (BioTek, Winooski, VT). Antibody data were analyzed by nonlinear regression (One site total binding) to determine EC_50_ titers in GraphPad Prism 8 (GraphPad Software, La Jolla, CA).

### Statistical Analysis

All statistical analysis was performed using GraphPad Prism 8 Software (GraphPad Software, La Jolla, CA). A *P*-value<0.05 was considered significant.

## Author Contributions

A.T.W designed and executed experiments, analyzed data, secured funding, and wrote the paper. H.F., G.G., and W.R. designed and executed experiments, analyzed data, and provided subject matter expertise. T.L., H.S., and K.V. generated data. M.K.M. analyzed data and provided subject matter expertise. S.F., A.S., and D.E. provided data on natural infection samples. T.E., S.T., and A.L.R. secured funding and provided subject matter expertise. R.G.J. provided project oversight, secured funding, and provided subject matter expertise. J.R.C designed and executed experiments, analyzed data, and secured funding.

## Data availability

The authors declare that all data supporting the findings of this study are available within this article and its Supplementary Information files, or from the corresponding author upon reasonable request. Single-cell RNAseq gene expression data have been deposited in the Gene Expression Omnibus database (GSE154386).

## Disclaimer

The opinions or assertions contained herein are the private views of the authors and are not to be construed as reflecting the official views of the US Army or the US Department of Defense, or the National Institutes of Health. Material has been reviewed by the Walter Reed Army Institute of Research. There is no objection to its presentation and/or publication. The investigators have adhered to the policies for protection of human subjects as prescribed in AR 70–25.

## LIST OF SUPPLEMENTAL MATERIAL

**Supplemental Figure 1.** DENV-specific IgM/IgG titers and duration of fever in the natural DENV-1 infection subjects included in this analysis

**Supplemental Figure 2.** Flow cytometry gating scheme for monocyte, DC, and B cell phenotyping from DHIM study samples

**Supplemental Figure 3.** Flow cytometry gating scheme for T and NK cell phenotyping for DHIM study samples

**Supplemental Figure 4.** Correlation between population frequencies as defined by scRNAseq or flow cytometry in all DHIM study samples. Population frequency defined as percent of all cells

**Supplemental Figure 5.** Plasmablast expansion following experimental DENV-1 infection in all DHIM study subjects

**Supplemental Figure 6.** Surface Ig expression and isotype distribution on naïve B cells and memory B cells from DHIM study days 14/15

**Supplemental Figure 7.** CD4+ T cell and CD8+ T cell activation in all analyzed DHIM study participants

**Supplemental Figure 8.** NK and NKT cell activation in all analyzed DHIM study participants

**Supplemental Figure 9.** Number of differentially express genes in the indicated populations relative to baseline

**Supplemental Figure 10.** Proposed model

**Supplemental Table 1.** Natural primary DENV infection **s**ubject information

**Supplemental Table 2.** Sample sequencing metrics

**Supplemental Table 3.** Sample/population frequency: T cell populations

**Supplemental Table 4.** Sample/population frequency: B cells and myeloid linage cells

**Supplemental Table 5.** mAb sequence information

**Supplemental Table 6**. mAb EC50 from DHIM subject 002, day 15 post infection

**Supplemental Table 7.** Conserved differentially expressed genes: day 10 DHIM

**Supplemental Table 8.** Core differentially expressed genes: natural DENV-1 infection

**Supplemental Table 9**. Antibodies used for flow cytometry

## REFERENCES

1. D. J. Gubler, Aedes aegypti and Aedes aegypti-borne disease control in the 1990s: top down or bottom up. Charles Franklin Craig Lecture. Am J Trop Med Hyg 40, 571–578 (1989).

2. S. Bhatt et al., The global distribution and burden of dengue. Nature 496, 504–507 (2013).

3. D. S. Shepard, L. Coudeville, Y. A. Halasa, B. Zambrano, G. H. Dayan, Economic impact of dengue illness in the Americas. Am J Trop Med Hyg 84, 200–207 (2011).

4. M. G. Guzman, E. Harris, Dengue. Lancet 385, 453–465 (2015).

5. N. Sangkawibha et al., Risk factors in dengue shock syndrome: a prospective epidemiologic study in Rayong, Thailand. I. The 1980 outbreak. Am J Epidemiol 120, 653–669 (1984).

6. S. Thein et al., Risk factors in dengue shock syndrome. Am J Trop Med Hyg 56, 566–572 (1997).

7. I. K. Yoon et al., Characteristics of mild dengue virus infection in Thai children. Am J Trop Med Hyg 89, 1081–1087 (2013).

8. M. Chan, M. A. Johansson, The incubation periods of Dengue viruses. PLoS One 7, e50972 (2012).

9. B. D. Kirkpatrick et al., The live attenuated dengue vaccine TV003 elicits complete protection against dengue in a human challenge model. Sci Transl Med 8, 330ra336 (2016).

10. T. P. Endy et al., A Phase 1, Open Label-Assessment of a Dengue Virus-1 Live Virus Human Challenge - (DENV-1-LVHC) Strain. J Infect Dis, (2020).

11. C. P. Larsen, S. S. Whitehead, A. P. Durbin, Dengue human infection models to advance dengue vaccine development. Vaccine 33, 7075–7082 (2015).

12. S. Green et al., Early immune activation in acute dengue illness is related to development of plasma leakage and disease severity. J Infect Dis 179, 755–762 (1999).

13. K. Haltaufderhyde et al., Activation of Peripheral T Follicular Helper Cells During Acute Dengue Virus Infection. J Infect Dis 218, 1675–1685 (2018).

14. A. T. Waickman et al., Transcriptional and clonal characterization of B cell plasmablast diversity following primary and secondary natural DENV infection. EBioMedicine 54, 102733 (2020).

15. U. K. Nivarthi et al., Longitudinal analysis of acute and convalescent B cell responses in a human primary dengue serotype 2 infection model. EBioMedicine 41, 465–478 (2019).

16. J. Wrammert et al., Rapid and massive virus-specific plasmablast responses during acute dengue virus infection in humans. J Virol 86, 2911–2918 (2012).

17. T. Balakrishnan et al., Dengue virus activates polyreactive, natural IgG B cells after primary and secondary infection. PLoS One 6, e29430 (2011).

18. J. W. Schoggins et al., A diverse range of gene products are effectors of the type I interferon antiviral response. Nature 472, 481–485 (2011).

19. J. W. Schoggins, C. M. Rice, Interferon-stimulated genes and their antiviral effector functions. Curr Opin Virol 1, 519–525 (2011).

20. W. M. Schneider, M. D. Chevillotte, C. M. Rice, Interferon-stimulated genes: a complex web of host defenses. Annu Rev Immunol 32, 513–545 (2014).

21. S. B. Halstead, E. J. O’Rourke, A. C. Allison, Dengue viruses and mononuclear phagocytes. II. Identity of blood and tissue leukocytes supporting in vitro infection. J Exp Med 146, 218–229 (1977).

22. Z. Kou et al., Monocytes, but not T or B cells, are the principal target cells for dengue virus (DV) infection among human peripheral blood mononuclear cells. J Med Virol 80, 134–146 (2008).

23. A. Srikiatkhachorn et al., Dengue viral RNA levels in peripheral blood mononuclear cells are associated with disease severity and preexisting dengue immune status. PLoS One 7, e51335 (2012).

24. M. O. Baclig et al., Flow cytometric analysis of dengue virus-infected cells in peripheral blood. Southeast Asian J Trop Med Public Health 41, 1352–1358 (2010).

25. A. D. King et al., B cells are the principal circulating mononuclear cells infected by dengue virus. Southeast Asian J Trop Med Public Health 30, 718–728 (1999).

26. F. Zanini, S. Y. Pu, E. Bekerman, S. Einav, S. R. Quake, Single-cell transcriptional dynamics of flavivirus infection. Elife 7, (2018).

27. F. Zanini et al., Virus-inclusive single-cell RNA sequencing reveals the molecular signature of progression to severe dengue. Proc Natl Acad Sci U S A 115, E12363–E12369 (2018).

28. M. A. Sanborn et al., Analysis of cell-associated DENV RNA by oligo(dT) primed 5’ capture scRNAseq. Sci Rep 10, 9047 (2020).

29. V. Vorndam, M. Beltran, Enzyme-linked immunosorbent assay-format microneutralization test for dengue viruses. Am J Trop Med Hyg 66, 208–212 (2002).

30. H. H. Houng, R. Chung-Ming Chen, D. W. Vaughn, N. Kanesa-thasan, Development of a fluorogenic RT-PCR system for quantitative identification of dengue virus serotypes 1-4 using conserved and serotype-specific 3’ noncoding sequences. J Virol Methods 95, 19–32. (2001).

31. H. H. Houng, D. Hritz, N. Kanesa-thasan, in International Congress of Virology. (Sydney, Australia, 1999).

32. M. Simmons, G. S. Murphy, T. Kochel, K. Raviprakash, C. G. Hayes, Characterization of antibody responses to combinations of a dengue-2 DNA and dengue-2 recombinant subunit vaccine. Am J Trop Med Hyg 65, 420–426 (2001).

33. D. W. Vaughn et al., Dengue in the early febrile phase: viremia and antibody responses. J Infect Dis 176, 322–330 (1997).

34. S. Kalayanarooj et al., Early clinical and laboratory indicators of acute dengue illness. J Infect Dis 176, 313–321 (1997).

35. D. H. Libraty et al., Differing influences of virus burden and immune activation on disease severity in secondary dengue-3 virus infections. J Infect Dis 185, 1213–1221 (2002).

36. B. L. Innis et al., An enzyme-linked immunosorbent assay to characterize dengue infections where dengue and Japanese encephalitis co-circulate. Am J Trop Med Hyg 40, 418–427 (1989).

37. G. X. Zheng et al., Massively parallel digital transcriptional profiling of single cells. Nat Commun 8, 14049 (2017).

38. T. Stuart et al., Comprehensive Integration of Single-Cell Data. Cell 177, 1888–1902 e1821 (2019).

39. A. Butler, P. Hoffman, P. Smibert, E. Papalexi, R. Satija, Integrating single-cell transcriptomic data across different conditions, technologies, and species. Nat Biotechnol 36, 411–420 (2018).

40. D. W. Lee et al., BRILIA: Integrated Tool for High-Throughput Annotation and Lineage Tree Assembly of B-Cell Repertoires. Front Immunol 7, 681 (2016).

