## Supplemental figures for "Temporally integrated single cell RNA sequencing analysis of controlled and natural primary human DENV-1 infections"

Supplemental Figure 1

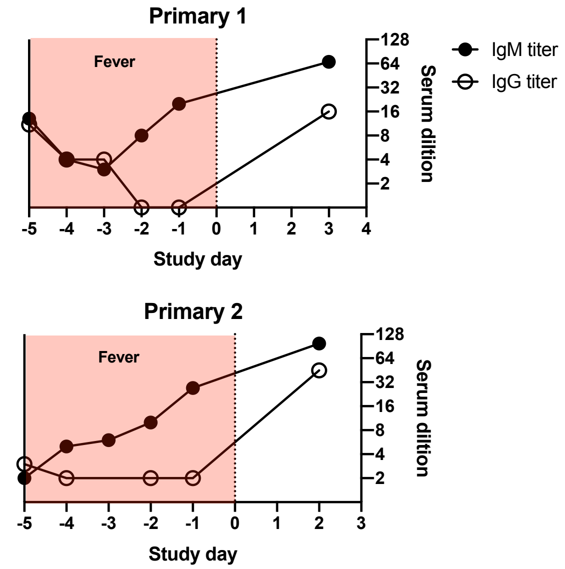


**Supplemental Figure 1.** DENV-specific IgM/IgG titers and duration of fever in the natural DENV-1 infection subjects included in this analysis

Supplemental Figure 2

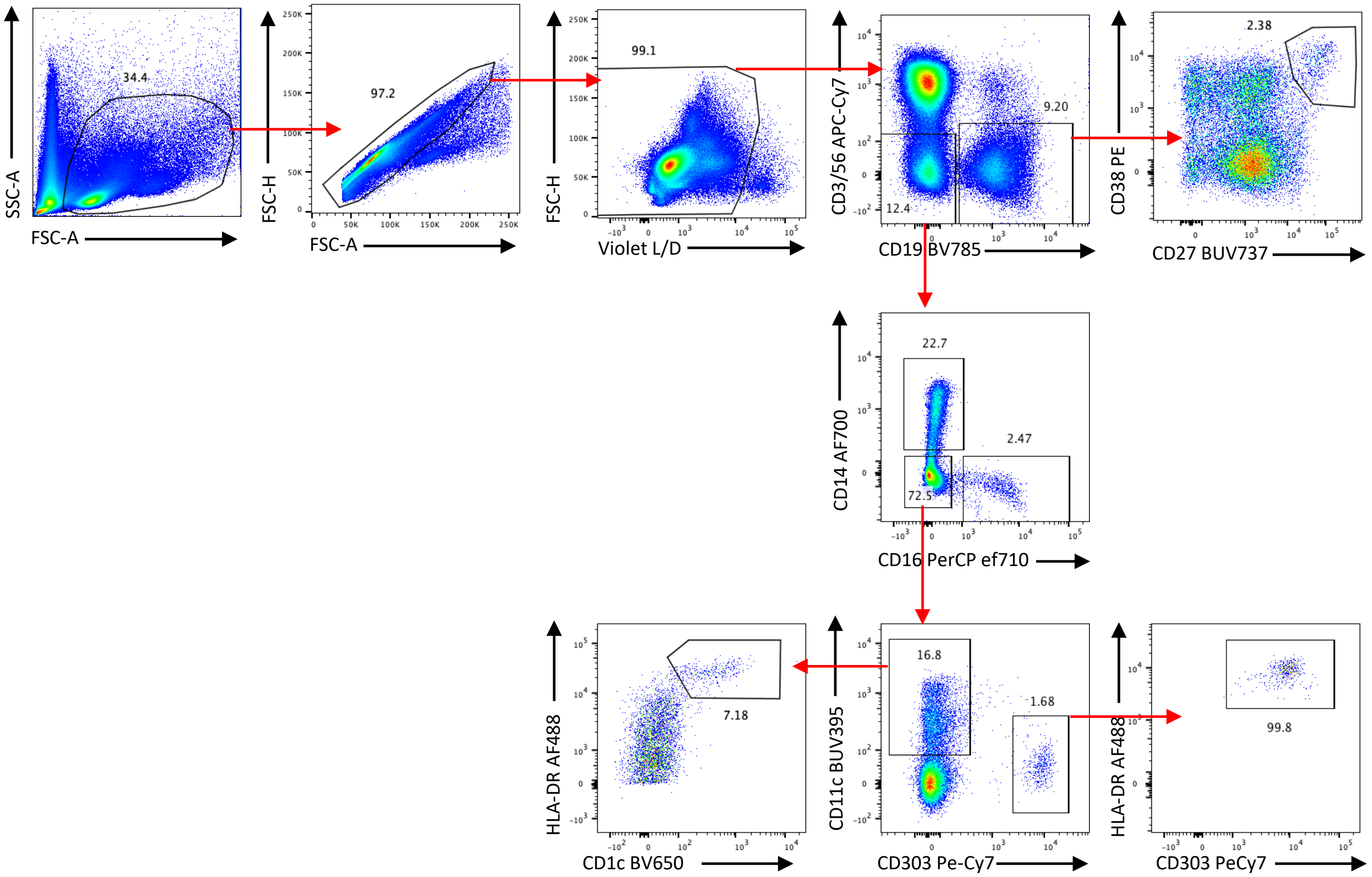


**Supplemental Figure 2.** Flow cytometry gating scheme for monocyte, DC, and B cell phenotyping from DHIM study samples

Supplemental Figure 3

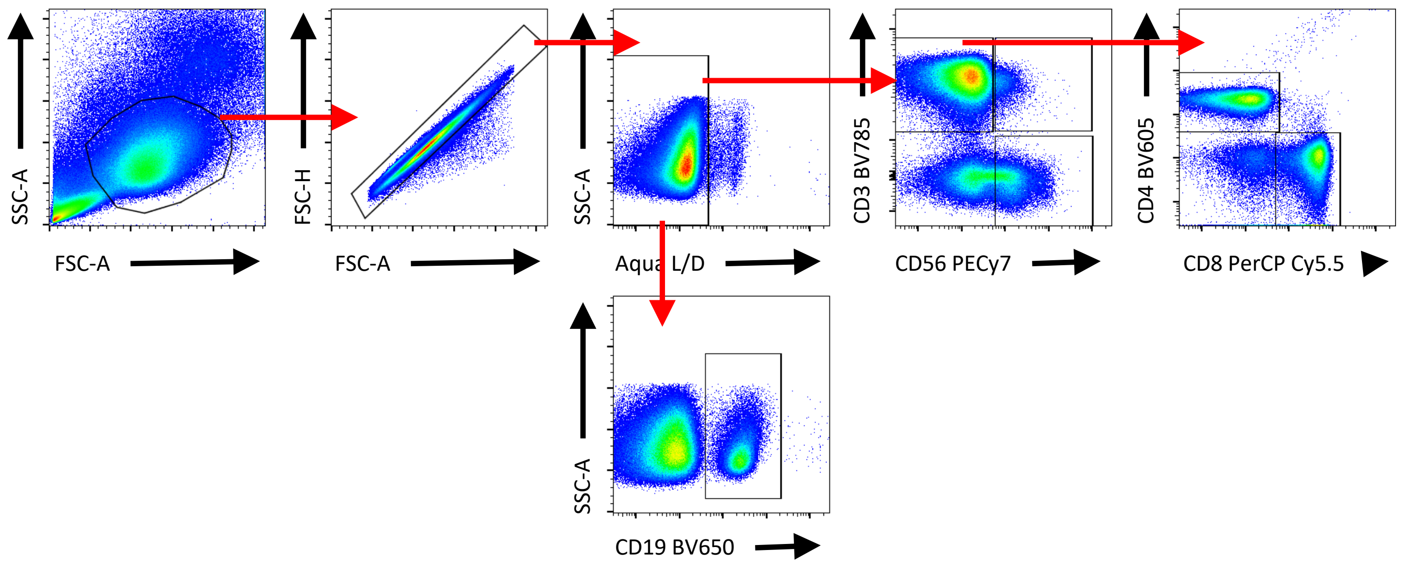


**Supplemental Figure 3.** Flow cytometry gating scheme for T and NK cell phenotyping for DHIM study samples

Supplemental Figure 4

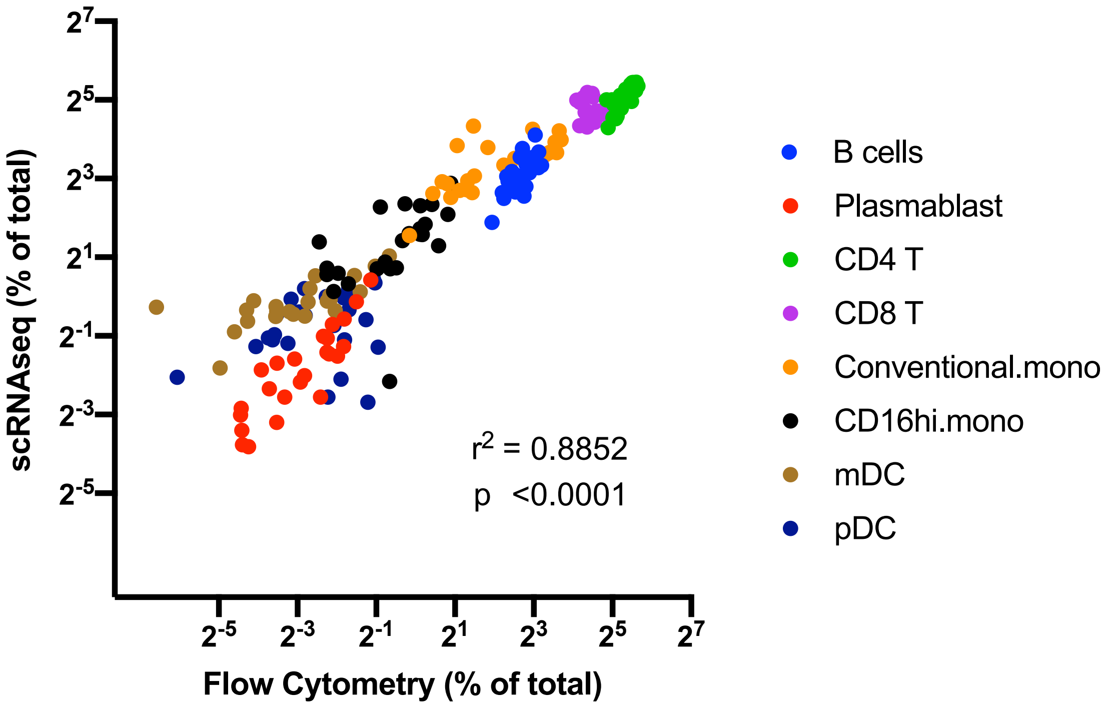


**Supplemental Figure 4.** Correlation between population frequencies as defined by scRNAseq or flow cytometry in all DHIM study samples. Population frequency defined as percent of all cells

Supplemental Figure 5

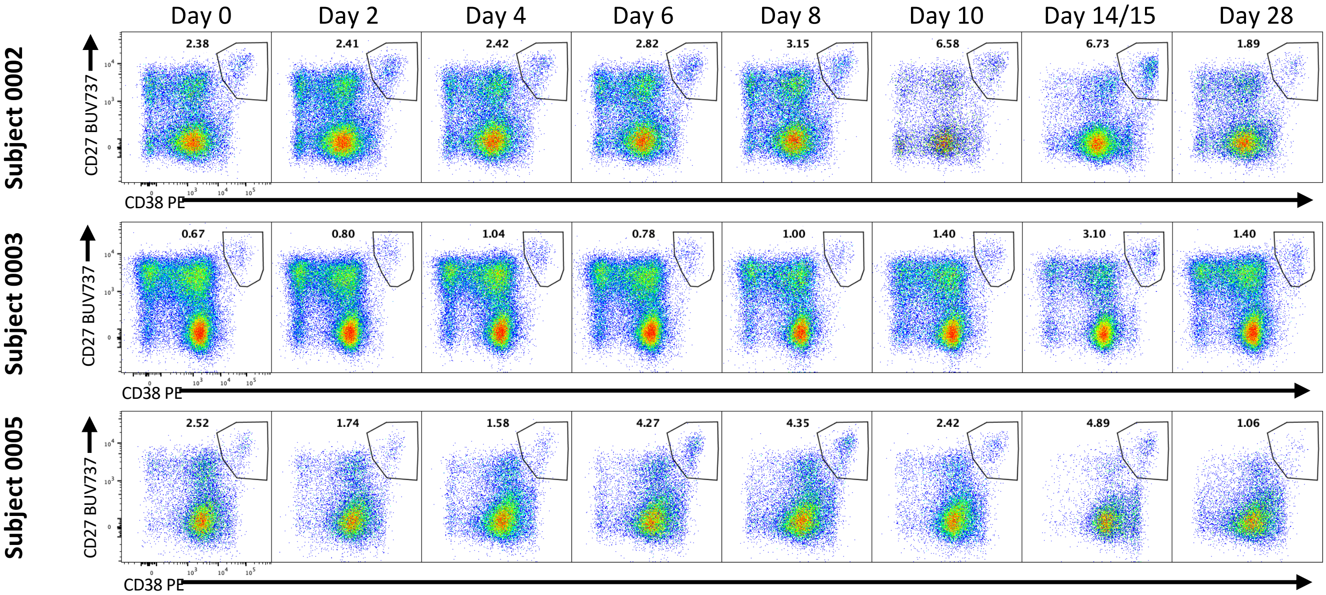


**Supplemental Figure 5.** Plasmablast expansion following experimental DENV-1 infection in all DHIM study subjects. Cells are gated on CD3-CD56-CD19+ viable lymphocytes

Supplemental Figure 6

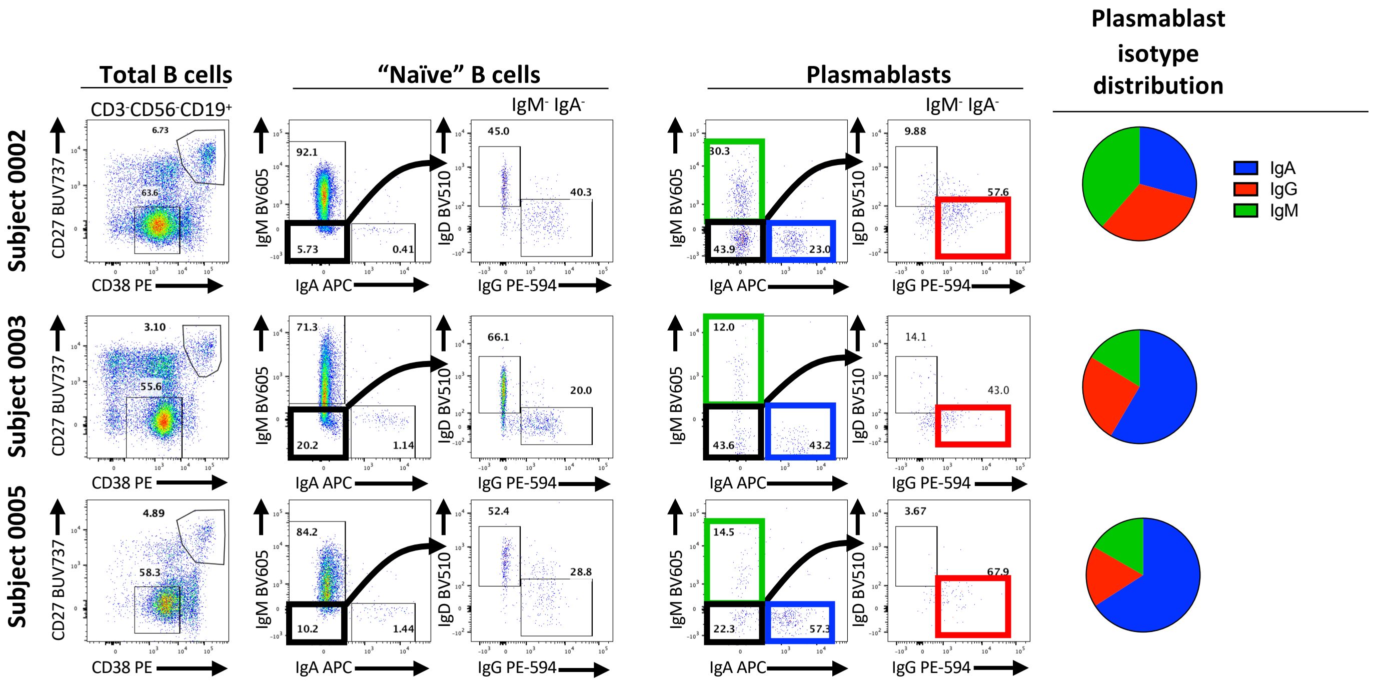


**Supplemental Figure 6.** Surface Ig expression and isotype distribution on naïve B cells and memory B cells from DHIM study days 14/15. Total B cells gated on viable CD3-CD56-CD19+ lymphocytes.

Supplemental Figure 7

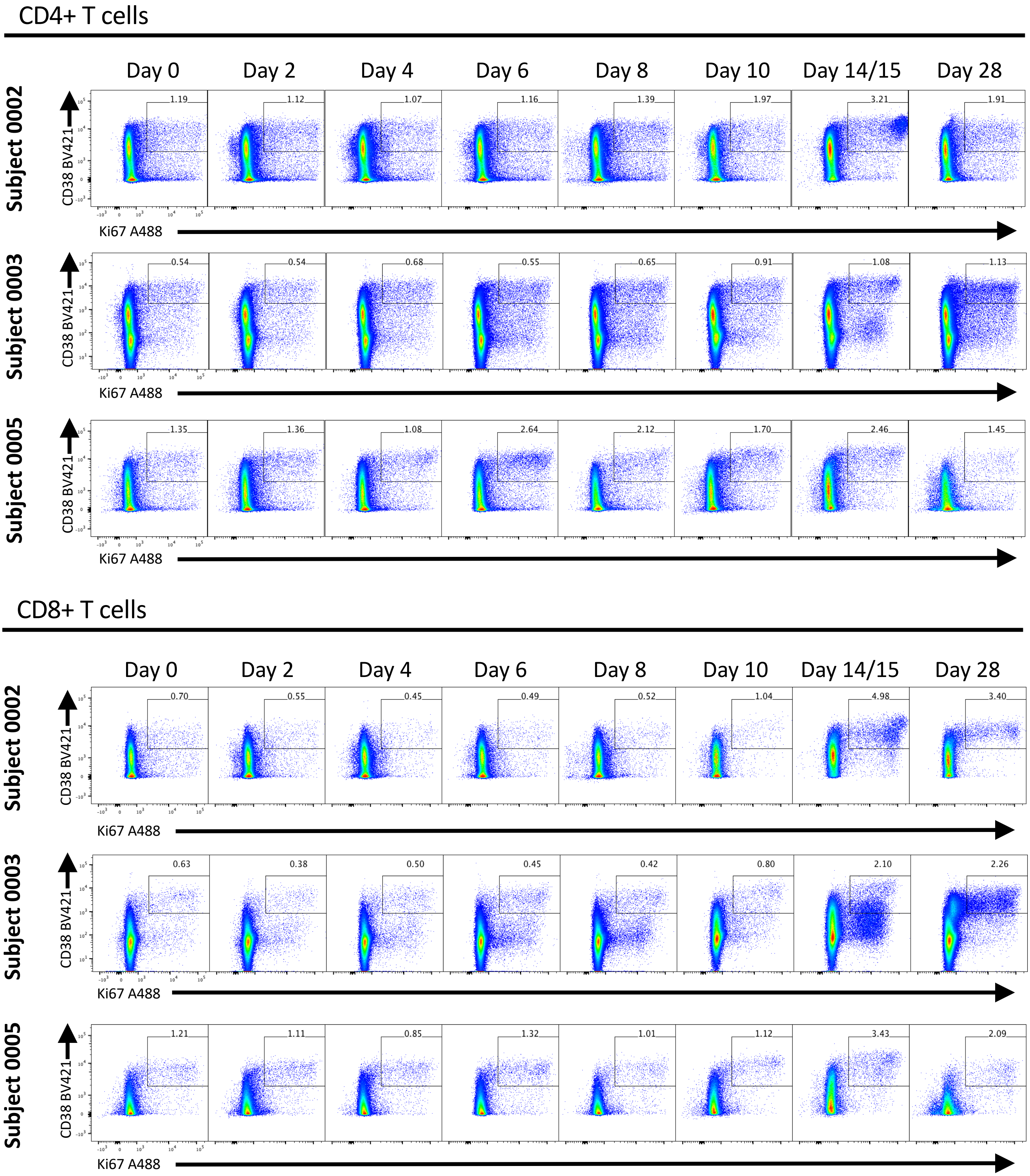


**Supplemental Figure 7.** CD4+ T cell and CD8+ T cell activation in all analyzed DHIM study participants. T cells defined as viable CD3+ CD56- lymphocytes.

Supplemental Figure 8

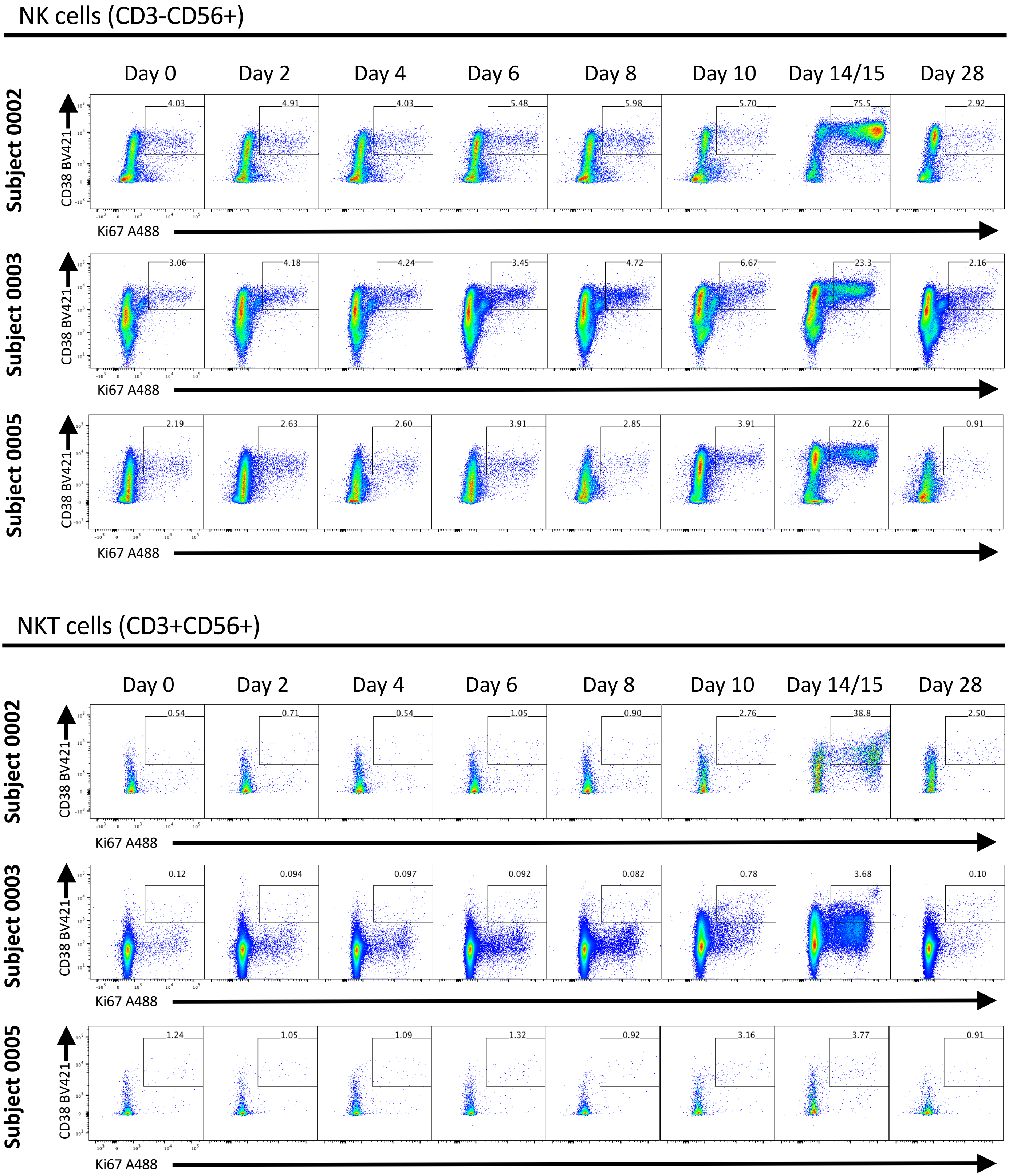


**Supplemental Figure 8.** NK and NKT cell activation in all analyzed DHIM study participants. NK cells defined as CD3- CD56+ viable lymphocytes. NKT cells defined as CD3+CD56+ viable lymphocytes

Supplemental Figure 9

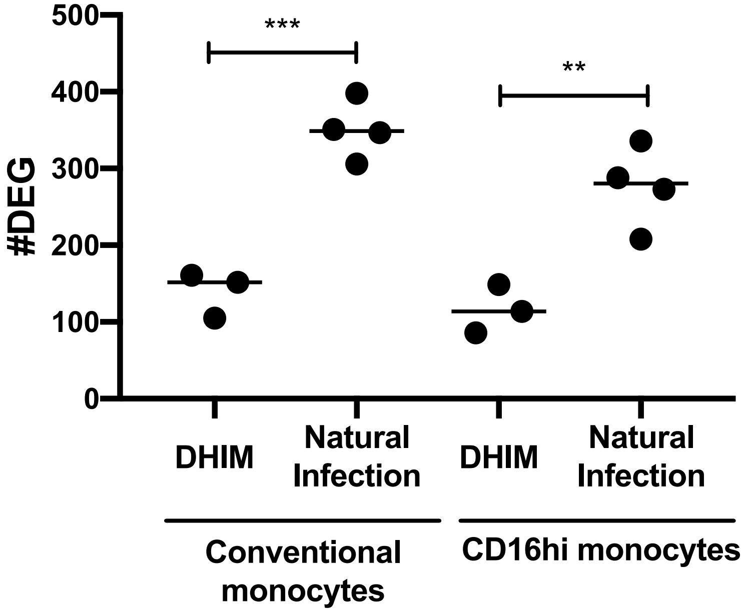


**Supplemental Figure 9.** Number of differentially express genes in the indicated populations relative to baseline. DHIM samples are from study day 10. Natural Infection samples are from acute 1/acute 2 time points. . ** p <0.01, *** p <0.001, unpaired two-tailed t test

Supplemental Figure 10

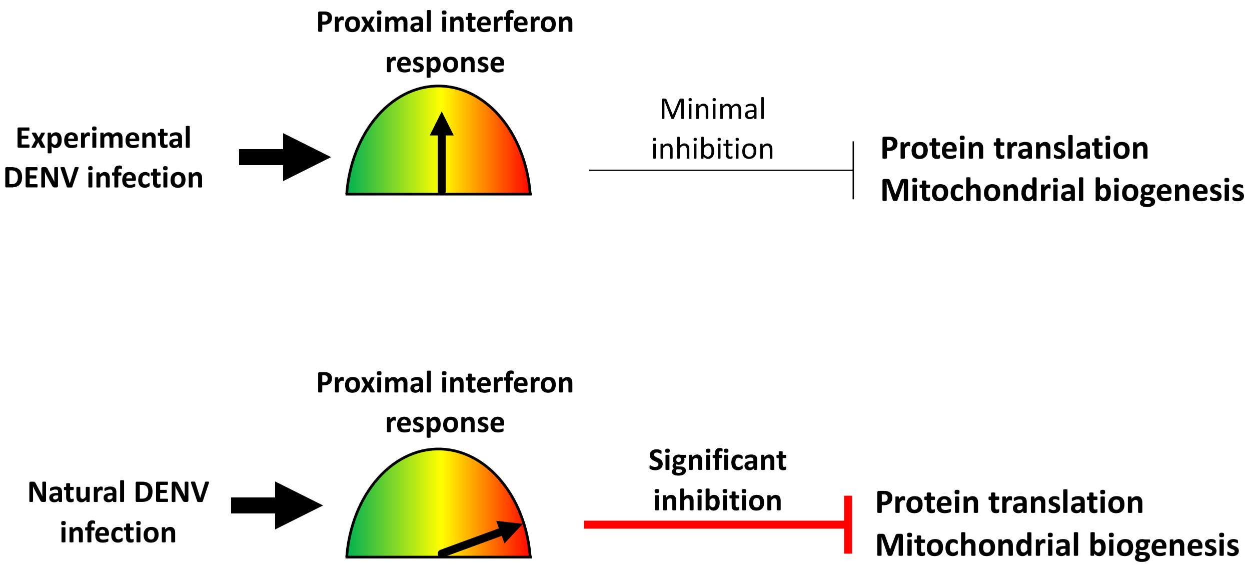


**Supplemental Figure 10**, Proposed model
