## Supplemental tables for "Temporally integrated single cell RNA sequencing analysis of controlled and natural primary human DENV-1 infections"

**Supplemental Table 1.** Natural primary DENV infection subject information

| Subject | Age / Sex | Infection | Serotype | Severity | Time point (fever day) |
|---|---|---|---|---|---|
| **Primary #1** | 9/M | Primary | DENV1 | DF | -3 |
| | | | | | -2 |
| | | | | | 180 |
| **Primary #2** | 5/M | Primary | DENV1 | DF | -5 |
| | | | | | -4 |
| | | | | | 180 |

**Supplemental Table 2.** Sample sequencing metrics

| Subject | Study day | Total reads | Recovered cells | Mean reads/cell | UMI/cell | Unique features/cell | Total features |
|---|---|---|---|---|---|---|---|
| DHIM #2 | 0 | 469,544,606 | 5,204 | 90,228 | 6,660 | 1,796 | 17,907 |
| DHIM #2 | 2 | 457,252,410 | 4,001 | 114,285 | 6,754 | 1,811 | 17,403 |
| DHIM #2 | 4 | 525,271,235 | 6,048 | 86,850 | 5,339 | 1,521 | 17,847 |
| DHIM #2 | 6 | 469,788,165 | 6,046 | 77,702 | 5,732 | 1,645 | 17,922 |
| DHIM #2 | 8 | 407,460,368 | 6,289 | 64,789 | 5,138 | 1,477 | 17,319 |
| DHIM #2 | 10 | 501,489,087 | 4,150 | 120,841 | 5,293 | 1,611 | 17,051 |
| DHIM #2 | 15 | 456,124,963 | 4,917 | 92,765 | 7,112 | 2,095 | 17,494 |
| DHIM #2 | 28 | 447,848,025 | 6,767 | 66,181 | 5,323 | 1,709 | 17,776 |
| DHIM #3 | 0 | 483,342,365 | 8,446 | 57,227 | 6,584 | 1,896 | 18,318 |
| DHIM #3 | 2 | 436,966,440 | 6,426 | 68,000 | 6,587 | 1,845 | 17,752 |
| DHIM #3 | 4 | 545,710,915 | 7,062 | 77,274 | 6,801 | 1,934 | 18,046 |
| DHIM #3 | 6 | 530,778,776 | 9,549 | 55,585 | 6,514 | 1,870 | 18,433 |
| DHIM #3 | 8 | 457,507,268 | 6,448 | 70,953 | 6,813 | 1,943 | 17,683 |
| DHIM #3 | 10 | 485,332,264 | 8,350 | 58,124 | 7,067 | 2,056 | 18,014 |
| DHIM #3 | 14 | 460,390,651 | 8,697 | 52,937 | 7,058 | 2,092 | 18,140 |
| DHIM #3 | 28 | 374,634,551 | 12,666 | 29,578 | 5,470 | 1,671 | 18,347 |
| DHIM #5 | 0 | 415,606,235 | 3,534 | 117,602 | 6,524 | 1,897 | 17,287 |
| DHIM #5 | 2 | 435,634,393 | 5,498 | 79,235 | 5,914 | 1,775 | 17,789 |
| DHIM #5 | 4 | 420,675,942 | 4,109 | 102,379 | 5,567 | 1,679 | 16,921 |
| DHIM #5 | 6 | 378,723,340 | 3,863 | 98,039 | 5,773 | 1,708 | 16,971 |
| DHIM #5 | 8 | 440,433,309 | 3,424 | 128,631 | 6,155 | 1,737 | 17,074 |
| DHIM #5 | 10 | 421,623,137 | 6,431 | 65,561 | 5,390 | 1,775 | 17,618 |
| DHIM #5 | 14 | 427,483,659 | 5,221 | 81,878 | 5,841 | 1,982 | 17,458 |
| DHIM #5 | 28 | 414,736,158 | 3,869 | 107,195 | 5,055 | 1,522 | 16,726 |
| Primary #1 | Acute 1 | 917,183,458 | 1,993 | 460,202 | 4,403 | 1,640 | 15,693 |
| Primary #1 | Acute 2 | 652,219,383 | 3,852 | 169,320 | 4,621 | 1,590 | 16,256 |
| Primary #1 | 180 | 704,437,542 | 5,292 | 133,114 | 5,113 | 1,523 | 17,201 |
| Primary #2 | Acute 1 | 695,123,220 | 3,613 | 192,395 | 3,336 | 1,279 | 15,573 |
| Primary #2 | Acute 2 | 682,392,037 | 3,196 | 213,514 | 1,793 | 876 | 14,896 |
| Primary #2 | 180 | 683,419,691 | 6,247 | 109,400 | 4,295 | 1,381 | 16,873 |

**Supplemental Table 3.** Sample/population frequency: T cell populations

| Subject | Study day | Naïve CD4 | Memory CD4 | CD4 T | Naïve CD8 | CD8 CM | CD8 EM | Treg | MAIT | Vd2 gdT | NK/NKT | ILC | Activ T |
|---|---|---|---|---|---|---|---|---|---|---|---|---|---|
| DHIM #2 | 0 | 1021 | 933 | 227 | 624 | 451 | 153 | 73 | 108 | 123 | 169 | 43 | 3 |
| DHIM #2 | 2 | 798 | 769 | 168 | 596 | 347 | 77 | 50 | 91 | 82 | 119 | 24 | 5 |
| DHIM #2 | 4 | 1317 | 1064 | 255 | 820 | 469 | 121 | 82 | 121 | 117 | 184 | 42 | 7 |
| DHIM #2 | 6 | 1106 | 1179 | 226 | 898 | 570 | 124 | 100 | 133 | 123 | 183 | 41 | 10 |
| DHIM #2 | 8 | 1172 | 1277 | 224 | 768 | 608 | 160 | 104 | 125 | 140 | 193 | 50 | 11 |
| DHIM #2 | 10 | 852 | 817 | 147 | 398 | 336 | 109 | 69 | 81 | 84 | 118 | 23 | 10 |
| DHIM #2 | 15 | 913 | 873 | 225 | 809 | 255 | 240 | 76 | 140 | 94 | 467 | 43 | 196 |
| DHIM #2 | 28 | 1261 | 1080 | 219 | 669 | 467 | 209 | 100 | 162 | 131 | 326 | 49 | 28 |
| DHIM #3 | 0 | 963 | 1373 | 298 | 241 | 1068 | 1273 | 59 | 125 | 255 | 260 | 35 | 17 |
| DHIM #3 | 2 | 745 | 825 | 164 | 156 | 699 | 1290 | 50 | 132 | 286 | 351 | 41 | 19 |
| DHIM #3 | 4 | 815 | 937 | 196 | 166 | 798 | 1364 | 43 | 133 | 326 | 363 | 45 | 20 |
| DHIM #3 | 6 | 949 | 1114 | 227 | 213 | 1071 | 1857 | 52 | 165 | 412 | 485 | 57 | 24 |
| DHIM #3 | 8 | 626 | 725 | 166 | 149 | 788 | 1417 | 43 | 98 | 351 | 380 | 31 | 14 |
| DHIM #3 | 10 | 895 | 839 | 187 | 169 | 809 | 1687 | 46 | 121 | 392 | 439 | 36 | 101 |
| DHIM #3 | 14 | 860 | 688 | 162 | 149 | 892 | 1892 | 45 | 99 | 485 | 551 | 36 | 121 |
| DHIM #3 | 28 | 1216 | 1435 | 312 | 295 | 1529 | 2710 | 86 | 228 | 584 | 564 | 93 | 43 |
| DHIM #5 | 0 | 443 | 436 | 113 | 289 | 267 | 206 | 38 | 107 | 61 | 168 | 26 | 10 |
| DHIM #5 | 2 | 799 | 668 | 160 | 502 | 447 | 493 | 61 | 170 | 98 | 404 | 42 | 6 |
| DHIM #5 | 4 | 718 | 713 | 156 | 537 | 442 | 108 | 59 | 142 | 45 | 159 | 16 | 12 |
| DHIM #5 | 6 | 585 | 621 | 133 | 447 | 414 | 141 | 63 | 188 | 83 | 186 | 27 | 8 |
| DHIM #5 | 8 | 433 | 570 | 92 | 375 | 374 | 100 | 58 | 227 | 102 | 131 | 16 | 11 |
| DHIM #5 | 10 | 959 | 763 | 172 | 551 | 445 | 403 | 69 | 167 | 63 | 432 | 32 | 17 |
| DHIM #5 | 14 | 837 | 582 | 183 | 395 | 341 | 607 | 53 | 170 | 109 | 583 | 49 | 99 |
| DHIM #5 | 28 | 475 | 654 | 111 | 331 | 399 | 141 | 67 | 245 | 110 | 157 | 19 | 18 |
| Primary #1 | Acute 1 | 308 | 374 | 89 | 122 | 171 | 123 | 34 | 36 | 50 | 36 | 13 | 52 |
| Primary #1 | Acute 2 | 521 | 561 | 136 | 306 | 289 | 351 | 35 | 55 | 123 | 185 | 16 | 88 |
| Primary #1 | 180 | 747 | 588 | 159 | 301 | 355 | 308 | 71 | 94 | 81 | 191 | 31 | 63 |
| Primary #2 | Acute 1 | 83 | 140 | 32 | 51 | 104 | 254 | 22 | 41 | 44 | 162 | 15 | 29 |
| Primary #2 | Acute 2 | 258 | 342 | 65 | 99 | 434 | 591 | 23 | 64 | 79 | 287 | 18 | 24 |
| Primary #2 | 180 | 1111 | 583 | 252 | 471 | 495 | 855 | 45 | 67 | 120 | 344 | 35 | 30 |

**Supplemental Table 4.** Sample/population frequency: B cells and myeloid linage cells

| Subject | Study day | Naïve B | Memory B | Plasmablast | Conventional mono | CD16hi mono | mDC | pDC | Mono platelet | Mega | Neutrophil | doublet |
|---|---|---|---|---|---|---|---|---|---|---|---|---|
| DHIM #2 | 0 | 316 | 210 | 19 | 435 | 65 | 40 | 52 | 29 | 81 | 9 | 20 |
| DHIM #2 | 2 | 224 | 165 | 15 | 263 | 66 | 38 | 51 | 16 | 23 | 4 | 10 |
| DHIM #2 | 4 | 429 | 346 | 29 | 348 | 66 | 55 | 59 | 27 | 71 | 14 | 5 |
| DHIM #2 | 6 | 346 | 281 | 30 | 392 | 89 | 56 | 69 | 26 | 40 | 15 | 9 |
| DHIM #2 | 8 | 408 | 328 | 22 | 393 | 95 | 49 | 42 | 21 | 69 | 17 | 13 |
| DHIM #2 | 10 | 217 | 125 | 38 | 421 | 109 | 32 | 57 | 34 | 58 | 13 | 2 |
| DHIM #2 | 15 | 181 | 107 | 66 | 144 | 11 | 14 | 23 | 6 | 27 | 3 | 4 |
| DHIM #2 | 28 | 298 | 173 | 15 | 967 | 328 | 63 | 78 | 25 | 99 | 8 | 12 |
| DHIM #3 | 0 | 310 | 436 | 8 | 1067 | 227 | 97 | 35 | 146 | 106 | 14 | 33 |
| DHIM #3 | 2 | 192 | 213 | 9 | 818 | 192 | 47 | 46 | 79 | 42 | 19 | 11 |
| DHIM #3 | 4 | 238 | 302 | 5 | 876 | 212 | 50 | 31 | 65 | 47 | 14 | 16 |
| DHIM #3 | 6 | 330 | 395 | 7 | 1458 | 342 | 80 | 45 | 91 | 116 | 27 | 32 |
| DHIM #3 | 8 | 187 | 217 | 8 | 781 | 213 | 51 | 33 | 76 | 67 | 12 | 15 |
| DHIM #3 | 10 | 227 | 245 | 23 | 1324 | 614 | 59 | 39 | 39 | 38 | 10 | 11 |
| DHIM #3 | 14 | 178 | 143 | 29 | 1618 | 447 | 72 | 21 | 91 | 86 | 16 | 16 |
| DHIM #3 | 28 | 481 | 669 | 25 | 1512 | 385 | 82 | 61 | 108 | 169 | 21 | 58 |
| DHIM #5 | 0 | 202 | 111 | 6 | 712 | 175 | 51 | 27 | 46 | 27 | 3 | 10 |
| DHIM #5 | 2 | 325 | 134 | 6 | 760 | 234 | 56 | 33 | 51 | 22 | 14 | 13 |
| DHIM #5 | 4 | 355 | 125 | 7 | 311 | 67 | 70 | 7 | 36 | 15 | 6 | 3 |
| DHIM #5 | 6 | 344 | 148 | 16 | 283 | 64 | 42 | 9 | 41 | 13 | 3 | 4 |
| DHIM #5 | 8 | 329 | 137 | 23 | 263 | 63 | 70 | 14 | 16 | 18 | 0 | 2 |
| DHIM #5 | 10 | 405 | 143 | 16 | 1225 | 327 | 65 | 51 | 63 | 26 | 10 | 27 |
| DHIM #5 | 14 | 253 | 63 | 32 | 595 | 128 | 28 | 38 | 25 | 43 | 4 | 4 |
| DHIM #5 | 28 | 500 | 167 | 12 | 238 | 63 | 56 | 6 | 15 | 73 | 4 | 8 |
| Primary #1 | Acute 1 | 236 | 133 | 92 | 73 | 20 | 6 | 10 | 0 | 12 | 0 | 3 |
| Primary #1 | Acute 2 | 328 | 354 | 164 | 141 | 34 | 10 | 115 | 5 | 29 | 2 | 4 |
| Primary #1 | 180 | 503 | 561 | 116 | 781 | 115 | 93 | 67 | 23 | 21 | 7 | 16 |
| Primary #2 | Acute 1 | 154 | 165 | 100 | 1664 | 389 | 35 | 80 | 33 | 13 | 1 | 2 |
| Primary #2 | Acute 2 | 153 | 109 | 51 | 331 | 159 | 7 | 51 | 14 | 35 | 0 | 2 |
| Primary #2 | 180 | 663 | 611 | 29 | 260 | 121 | 41 | 41 | 38 | 24 | 2 | 9 |

**Supplemental Table 5.** mAb sequence information

| Clone name | Subject | Parental Isotype | Heavy chain CDR3aa | Heavy chain V | Heavy chain D | Heavy chain J | Light chain CDR3aa | Light chain V | Light chain J |
|---|---|---|---|---|---|---|---|---|---|
| **VDB-63** | DHIM-0002 | IgM | CARDPGGTIANNCFDSW | IGHV3-33 | IGHD3-16 | IGHJ5 | CQQGYTSPLTF | IGKV4-1 | IGKJ4 |
| **VDB-64** | DHIM-0002 | IgM | CARRIAAAGRVDYW | IGHV5-51 | IGHD6-13 | IGHJ4 | CQQYGSSPSWTF | IGKV3-20 | IGKJ1 |
| **VDB-65** | DHIM-0002 | IgM | CARRDYDTSGSDYW | IGHV5-51 | IGHD3-22 | IGHJ4 | CQSADSSGTWVF | IGLV3-25 | IGLJ3 |
| **VDB-66** | DHIM-0002 | IgM | CARQSRGSSWYGSPFDVW | IGHV4-39 | IGHD6-13 | IGHJ3 | CGTWDSSLSAEVF | IGLV1-51 | IGLJ3 |
| **VDB-67** | DHIM-0002 | IgM | CARDPGWGALDIW | IGHV3-7 | IGHD7-27 | IGHJ3 | CMQGTHGPYTF | IGKV2-30 | IGKJ2 |
| **VDB-68** | DHIM-0002 | IgG1 | CVTDLYHIDYFFDYW | IGHV1-24 | IGHD2-2 | IGHJ4 | CYSAADNKGVF | IGLV3-27 | IGLJ3 |
| **VDB-69** | DHIM-0002 | IgG1 | CARVDWNHVYYMDVW | IGHV4-59 | IGHD1-14 | IGHJ6 | CSSYTITSIVVF | IGLV2-14 | IGLJ2 |
| **VDB-70** | DHIM-0002 | IgG1 | CARGPSGRYFQHFQHW | IGHV3-7 | IGHD1-26 | IGHJ1 | CQSYDSSLSGWVF | IGLV1-40 | IGLJ3 |
| **VDB-71** | DHIM-0002 | IgG1 | CARARIKIFEVVDREFDYW | IGHV1-2 | IGHD3-3 | IGHJ4 | CQSYDIRLSGSVF | IGLV1-40 | IGLJ2 |
| **VDB-72** | DHIM-0002 | IgG2 | CARVPGHLW | IGHV3-53 | IGHD3-9 | IGHJ4 | CQEYNSYSLTF | IGKV1-5 | IGKJ4 |
| **VDB-73** | DHIM-0002 | IgA1 | CARGLGHNARYYYMDVW | IGHV4-61 | IGHD2-8 | IGHJ6 | CTTWDDNLNGPVF | IGLV1-44 | IGLJ2 |
| **VDB-74** | DHIM-0002 | IgA1 | CVRGRKTGGYDCYDYW | IGHV1-2 | IGHD5-12 | IGHJ4 | CQQYNSPITF | IGKV1-5 | IGKJ5 |
| **VDB-75** | DHIM-0002 | IgA1 | CARDSKEYNWNYADYFDFW | IGHV4-61 | IGHD1-7 | IGHJ4 | CQQYGSSPSTF | IGKV3-20 | IGKJ4 |
| **VDB-76** | DHIM-0002 | IgA1 | CATGGLSGDIRGIDFW | IGHV1-24 | IGHD7-27 | IGHJ4 | CYSAADNSRGVF | IGLV3-27 | IGLJ3 |
| **VDB-77** | DHIM-0002 | IgA1 | CARWLQLWESGWSFDLW | IGHV4-34 | IGHD5-18 | IGHJ2 | CQQYYDTPYTF | IGKV4-1 | IGKJ2 |

**Supplemental Table 6.** mAb EC50 from DHIM subject 002, day 15 post infection

| | | **mAb EC50 (ng/ml)** | | | |
|---|---|---|---|---|---|
| **Clone name** | **Isotype** | **DENV1** | **DENV2** | **DENV3** | **DENV4** |
| VDB-63 | IgM | >5,000 | >5,000 | >5,000 | >5,000 |
| VDB-64 | IgM | >5,000 | >5,000 | >5,000 | >5,000 |
| VDB-65 | IgM | >5,000 | >5,000 | >5,000 | >5,000 |
| VDB-66 | IgM | >5,000 | >5,000 | >5,000 | >5,000 |
| VDB-67 | IgM | >5,000 | >5,000 | >5,000 | >5,000 |
| VDB-68 | IgG1 | >5,000 | >5,000 | >5,000 | >5,000 |
| VDB-69 | IgG1 | 1,654 | >5,000 | >5,000 | >5,000 |
| VDB-70 | IgG1 | >5,000 | >5,000 | >5,000 | >5,000 |
| VDB-71 | IgG1 | >5,000 | >5,000 | >5,000 | >5,000 |
| VDB-72 | IgG2 | >5,000 | >5,000 | >5,000 | >5,000 |
| VDB-73 | IgA1 | 2,319 | 3,449 | 2,340 | 2,214 |
| VDB-74 | IgA1 | >5,000 | >5,000 | >5,000 | >5,000 |
| VDB-75 | IgA1 | >5,000 | >5,000 | >5,000 | >5,000 |
| VDB-76 | IgA1 | >5,000 | >5,000 | >5,000 | >5,000 |
| VDB-77 | IgA1 | >5,000 | >5,000 | >5,000 | >5,000 |

**Supplemental Table 7.** Conserved differentially expressed genes: day 10 DHIM

| **Population** | **Induced core genes** | **Suppressed core genes** |
|---|---|---|
| **Conventional monocytes** | IFITM1, IFI6, LY6E, IFITM3, ISG15, IFI44L, IFITM2, TYMP, XAF1, PSME2, RNF213, SIGLEC1, NCF1, TNFSF10, LAP3, APOBEC3A, MT2A, PLAC8, SERPING1, UBE2L6, TNFSF13B, IRF7, PSMB9, IFI35, PARP14, VAMP5, MX1, EPSTI1, TRIM22, EIF2AK2, OAS1, IFI44, RSAD2, GBP1, OAS3, HBB, WARS, STAT1, TMEM123, MX2, STAT2, PLSCR1, ISG20, SAMD4A, HES4, OASL, TXNIP | IL1B, CXCL8, RPS3A, RPL4, EEF2, RPL6, RPL5, EIF3L, RPL3, EIF4B |
| **CD16hi monocytes** | IFITM1, IFITM3, LY6E, IFI6, APOBEC3A, IFI44L, RSAD2, TNFSF10, TYMP, ISG15, BST2, UBE2L6, XAF1, LAP3, TNFSF13B, NCF1, PLAC8, IFI35, PSMB9, EPSTI1, IFIT2, TRIM22, PLSCR1, VAMP5, PSME2, SERPING1, IRF7, MX1, PARP14, EIF2AK2, WARS, IFI44, MX2, CXCL10, HES4, GBP1, TMEM123, OAS3, HERC5, OAS1 | MT-CYB, IL1B, RPL3, MT-ND4, RPL10A, AHNAK |

| | | |
|---|---|---|
| **mDC** | IFI6, IFITM1, LY6E, IFITM3, IFI44L, IFITM2, ISG15, UBE2L6, PSME2, TYMP, PLAC8, XAF1, MX1, STAT1, PSMB9, EPSTI1, LAP3, MT2A, IRF7, LGALS9, NAPA, MX2 | EEF1B2, EIF3L, RPL5 |
| **pDC** | LY6E, PLAC8, ISG15, BST2, IFI44L, IFITM2, COX5A, IFI35 | -- |
| **MAIT** | IFITM1, IFI6, LY6E, IFITM3, IFITM2, IFI44L, BST2, ISG20, ISG15, XAF1, PSME2, TRIM22, PSMB9, STAT1, IRF7, TYMP, IFI35, EPSTI1 | -- |
| **ILC** | IFITM1, IFI6, IFI44L, LY6E | -- |
| **NK/NKT** | IFI6, IFITM1, LY6E, IFI44L, XAF1, IFITM3, BST2, IFITM2, ISG15, PSME2, PLSCR1, IFI35, TRIM22, SHISA5, ISG20, MX2, EIF2AK2, UBE2L6, PSMB9, EPSTI1, MX1, IRF7, PARP9, TYMP, STAT1, LGALS1 | KLRB1 |
| **Vδ2 γδ T** | IFITM1, IFI6, LY6E, XAF1, ISG20, IFI44L, IFITM2, ISG15, STAT1, TRIM22, BST2, PLSCR1, EPSTI1, IFI35, MX2, EIF2AK2, PSME1, IRF7 | KLRB1 |
| **Naïve B** | IFITM1, IFI6, IFI44L, XAF1, LY6E, ISG15, EIF2AK2, IRF7, IFITM2, ISG20, TRIM22, BST2, MX2, RNF213, MX1, STAT1, PLSCR1, DRAP1, PSMB9, EPSTI1, PSME2 | -- |
| **Memory B** | IFI6, IFITM1, LY6E, ISG15, IFITM3, ISG20, XAF1, BST2, MX1, IFITM2, TRIM22, IFI44L, EIF2AK2, PSME2, STAT1, IRF7, MX2 | -- |
| **Naïve CD4** | IFITM1, IFI6, IFI44L, LY6E, HBB, STAT1, XAF1, TRIM22, ISG15, EIF2AK2, SP100, EPSTI1, PLSCR1, BST2, ISG20 | -- |
| **Memory CD4** | IFI6, IFITM1, LY6E, IFI44L, ISG15, XAF1, BST2, EIF2AK2, ISG20, TRIM22, STAT1, IFITM2, PSMB9, IFI44, EPSTI1, SP100, PARP9, PLSCR1, MX2 | -- |
| **Naïve CD8** | IFI6, IFITM1, IFI44L, LY6E, STAT1, TRIM22, BST2, ISG15, ISG20, XAF1, PSME2, IFI44, EIF2AK2, IFITM2 | -- |
| **CD8 CM** | IFI6, IFITM1, LY6E, IFI44L, XAF1, BST2, TRIM22, EIF2AK2, STAT1, ISG20, ISG15, SP100, PSMB9, EPSTI1, PLSCR1 | -- |
| **CD8 EM** | IFI6, IFITM1, LY6E, BST2, ISG15, IFI44L, PSME2, ISG20, STAT1, IFITM3, TRIM22, IFITM2, PLSCR1, IRF7, MX1, PSMB9, EIF2AK2, XAF1, IFI35, MT2A, SP100, DRAP1, MX2 | -- |
| **Treg** | IFI6, IFITM1, LY6E, ISG15, SP100, TRIM22, EIF2AK2, DRAP1 | RPL4 |

**Supplemental Table 8.** Core differentially expressed genes: natural DENV-1 infection

| Population | Induced core genes | Suppressed core genes |
|---|---|---|
| **Conventional monocytes** | IFIT3, IFI27, IFIT1, RSAD2, USP18, IFI44L, OASL, SERPING1, CCL2, OAS3, IFIT2, IFITM1, MX1, HERC5, ISG15, ZBP1, GBP1, SIGLEC1, XAF1, CMPK2, IFI6, LGALS3BP, SAMD9L, ISG20, PARP9, OAS2, EPSTI1, IFI44, IFIH1, STAT1, LY6E, SPATS2L, CXCL10, IRF7, OAS1, HLA-A, PARP12, EIF2AK2, PARP14, DDX58, TNFSF10, MX2, DDX60L, RNF213, GBP5, GBP4, TRIM22, SAMD9, PLAC8, CTSL, IFI35, MT2A, IFI16, NT5C3A, HLA-B, APOBEC3A, HLA-C, TCN2, IFITM2, HSH2D, LAP3, SP110, GIMAP4, IFITM3, SAMD4A, XRN1, SELL, RNASE2, NMI, FCGR1A, BST2, SMCHD1, PML, PSME2, PSMB9, PLSCR1, UBE2L6, GCH1, VAMP5, PHF11, TMEM123, TYMP, DRAP1, NAPA, SCO2, CCR1, LGALS9, IL1RN, PSMA4, MYL12A, MARCKS, MAFB | TPT1, RPS28, EEF1A1, RPS13, RPS3A, RPLP1, RPL8, RPL18, RPS7, RPS14, RPL34, RPS24, RPL35A, RPS15A, MT-ND3, MT-CO3, RPS8, RPS4X, RPL11, RPL10, RPL19, RPL5, RPL6, RPL9, NACA, RPL22, RPL26, RPL37, RPS23, RPL15, RPL7A, RPL32, RPL18A, BTF3, RPL3, RPL30, RPLP2, RPS16, RPS27A, RPS6, RPLP0, RPL29, COX4I1, PABPC1, CSTA, COTL1, SLC25A6, RPL4, RPL27, RPL13, RPL10A, RPS5, EEF1B2, RPL21, AP1S2, RPS3, EEF2, GSTP1, RPS18, PCBP2, RPL7, EIF3L, GNB2L1, SLC25A5, RPS4Y1, ALDH2, EIF3F, RBM3, FCGRT, EIF3E, EIF3H, CRTAP |
| **CD16hi monocytes** | IFI44L, IFIT1, OASL, RSAD2, SIGLEC1, IFI27, IFIT3, MX1, IFI44, IFIT2, CXCL10, XAF1, HERC5, LGALS3BP, ISG15, SERPING1, DDX60L, OAS3, PLSCR1, USP18, MX2, APOBEC3A, EIF2AK2, TNFSF10, IRF7, FFAR2, CD300E, ZBP1, EPSTI1, GBP4, NCF1, TCN2, GLUL, IFI35, DDX58, SAMD9L, LGALS9, HLA-A, IFIH1, SPATS2L, TRIM22, IFI6, MARCKS, CDKN1A | RPL6, RPS3A, EEF1A1, RPS15A, RPL10, RPS13, RPS8, RPS23, RPL34, RPS4X, RPS27A, RPL8, RPL14, RPS14, RPS7, RPL18A, RPS6, RPL12, RPLP2, RPL35A, RPL18, RPL11, RPL21, RPL7A, RPL9, RPL5, SLC25A6, RPL32, RPL10A, RPS24, RPS21, RPL13, RPL30, RPL4, MT-CO3, RPL19, RPL22, RPS16, RPL26, RPL15, GNB2L1, |

| | | |
|---|---|---|
| | | COTL1, PABPC1, RPL3, SLC25A5, ACTG1, EEF2, EEF1B2, NACA, RPLP0, RPL27, RPS25, RPS5, TPT1, RPL23A, RPS3, RPSA, RPS18, PFDN5, EIF3K, RPL37A, UQCRB, BTF3, RPL29, MT-ND3, MT-CYB, RPS4Y1, RPL23, YBX1, CCNI, RPS17, MT-ND4, NAP1L1, EIF3L, RBM3 |
| **mDC** | IFI27, USP18, IFIT3, RSAD2, HERC5, IFI44L | -- |
| **pDC** | IFI6, IFI44L, IFITM1, ISG15, ISG20, MX1, IFITM2 | -- |
| **MAIT** | IFI44L, IFI6, MX1, ISG15, MX2, IRF7, LY6E, IFITM1, IFI44, OAS1, IFIT1, IFIT3, MT2A, RSAD2, IFI35, OASL, PLSCR1, XAF1, ISG20, OAS3, BST2, TRIM22, UBE2L6, DRAP1, IFITM3 | RPS27A, EEF1A1, RPS4X, RPL3, RPS8, RPL13, MT-CYB, RPS3A, RPS14, RPS3, RPS6, MT-CO3, RPLP2, RPL10, MT-ND3, RPL21, MT-ATP6, RPL5, MT-ND5, RPL4 |
| **ILC** | IFI44L, IFI6, IFITM1 | -- |
| **NK/NKT** | IFI44L, IFIT3, MX1, IFI6, OAS1, OAS3, IFIT1, RSAD2, PLSCR1, IRF7, ISG15, OASL, MX2, EPSTI1, PARP9, ISG20, LY6E, IFI35, XAF1, IFI44, DTX3L, LGALS9, OAS2, MT2A, STAT1, TYMP, BST2, SAMD9L, EIF2AK2, ADAR, RNF213, SP100, SAMD9, GBP1, TRIM22, IFITM1, NT5C3A, CD38, PARP12, LAG3, UBE2L6, PRF1, ZBP1, S100A11, WARS, IFITM3, IFI16 | EEF1A1, RPS14, RPL13, RPL3, RPS3A, MT-CO3, RPS27A, MT-ATP6, RPS4X, RPL21, RPS8, MT-CYB, RPL5, MT-ATP8, KLRB1 |
| **Vδ2 γδ T** | IFI44L, IFI6, ISG15, LY6E, MX1, IFITM1, MT2A, IRF7, XAF1, IFIT3, RSAD2, IFITM3, OAS1, ISG20, CMPK2, STAT1, MX2, RNF213, UBE2L6, BST2, LAG3, SAMD9L, PLSCR1, SAMD9, IFITM2, TYMP, PSME2 | RPS27A, MT-CYB, RPS14, RPS3A, RPL5, RPL3, RPL21, EEF1A1, RPS3, RPS4X, RPLP2, RPS8, MT-CO3, EEF1B2, MT-ND4L, MT-ND5 |
| **Naïve B** | IFI44L, IFITM1, XAF1, ISG15, IFI6, IFIT3, MX1, IFIT1, MX2, IRF7, LY6E, PLSCR1, ISG20, HLA-A, CMPK2, EPSTI1, SAMD9L, OAS1, STAT1, UBE2L6, EIF2AK2, TRIM22, IFITM2, IFITM3, DRAP1, IFI44, HERC5, BST2, SP110, SAMD9, RNF213, IFI35, ZBP1, PARP9, PSME2, PSMB9, IFI16, MT2A, PLAC8, COX5A, STAT2, CHMP5 | RPS8, RPS3A, RPL3, RPS6, MT-CO3, RPL10A, RPL21, MT-CYB, MT-ATP6, RPL4, EIF3L |
| **Memory B** | IFIT3, IFI6, IFIT1, XAF1, ISG15, MX1, IRF7, IFI44L, MX2, IFITM1, HERC5, LY6E, STAT1, OAS1, PLSCR1, EIF2AK2, SAMD9L, ISG20, PARP9, EPSTI1, BST2, TRIM22, CTA-384D8.34, RSAD2, UBE2L6, MT2A, OAS2, RNF213, IFITM2, IFI35, ZBP1, SAMD9, IFITM3, DRAP1, PPM1K, DDX60L, TYMP, SAT1, CLEC2D, FCRL5 | RPS8, RPS3A, RPS6, RPS4X, RPL3, RPL21, RPL10A, RPL27, MT-CO3, MT-CYB, RPL4, EIF3L |
| **Naïve CD4** | IFI44L, ISG15, MX1, XAF1, IFIT3, LY6E, IRF7, HLA-A, IFI6, IFI44, OAS1, IFITM1, MX2, STAT1, EIF2AK2, PLSCR1, MT2A, EPSTI1, IFI35, BST2, TRIM22, RNF213, UBE2L6, DRAP1, IFI16, SMCHD1, MYL12A, C19orf66, ISG20, PSME2, PARP10, SAT1, IRF1 | RPS14, RPS3A, RPS6, RPL3, RPS27A, RPL21, RPL4, RPS16, MT-CYB, EIF3L |
| **Memory CD4** | IFI44L, ISG15, MX1, IFI6, IFIT3, XAF1, LY6E, IRF7, MX2, HLA-A, OAS1, IFITM1, MT2A, HLA-B, EPSTI1, HLA-E, STAT1, RSAD2, EIF2AK2, IFI44, ISG20, OASL, PLSCR1, SAMD9L, TRIM22, BST2, SAMD9, SP100, PARP9, RNF213, OAS3, HERC5, GBP1, IFI35, TYMP, SP110, UBE2L6, TNFSF10, IFI16, PHF11, IER2, DRAP1, ADAR, PSMB9, LGALS9, ZBP1, PARP12, PSME2, SMCHD1, PARP10, IFITM3 | EEF1A1, RPS3A, RPS27A, RPL13, RPS8, RPL34, RPS14, RPL3, RPS4X, RPS6, RPS18, RPL10, TPT1, RPL18, RPL21, RPL5, RPLP2, RPS5, RPL6, RPL10A, RPL4, RPL27, RPS16, MT-CYB, MT-CO3, EEF2, EIF3L |
| **Naïve CD8** | IFI44L, ISG15, MX1, XAF1, IFI6, IRF7, LY6E, IFIT3, IFI44, STAT1, IFITM1, EPSTI1, EIF2AK2, MT2A, OAS1, MX2, PLSCR1, BST2, SAMD9, IFI35, TRIM22, MYL12A, RSAD2, PARP10, | RPS6, RPS27A, RPS3A, RPL3, RPL21, MT-CO3, RPL4, RPS16, MT-CYB, MT-ATP6, EIF3L |

| | | |
|---|---|---|
| | RNF213, DRAP1, IFI16, SAMD9L, IFITM3, ISG20, PSMB9 | |
| **CD8 CM** | IFI44L, ISG15, MX1, IFI6, XAF1, LY6E, OAS1, IFITM1, IRF7, IFIT3, EIF2AK2, MX2, PTPRCAP, RSAD2, ISG20, EPSTI1, BST2, IFI35, STAT1, SP100, PLSCR1, MT2A, PARP9, OASL, RNF213, OAS2, TYMP, TRIM22, SAMD9, SAMD9L, DRAP1, IFI16, UBE2L6, IFITM3, PSME2, ZBP1, PSMB9, PARP10, LAG3 | RPS4X, MT-CYB, RPS3A, RPS27A, RPL3, EEF1A1, RPL13, RPS14, MT-CO3, RPL21, RPS6, RPS8, RPLP2, RPS16, RPL5, RPL4, MT-ATP6, MT-ND4L, MT-ND5, EIF3L, EEF2 |
| **CD8 EM** | MX1, IFI6, ISG15, IFI44L, XAF1, IFIT3, IFIT1, RSAD2, EIF2AK2, OAS1, MX2, ISG20, LY6E, IRF7, STAT1, MT2A, IFITM1, IFI35, PARP9, EPSTI1, BST2, IFI44, GBP1, PLSCR1, LAG3, UBE2L6, RNF213, TYMP, OASL, LGALS9, OAS2, SP100, PSME2, SAMD9L, TRIM22, HERC5, SP110, ADAR, TAP1, SAMD9, PSMB9, PRF1, IFI16, CD38, DRAP1, S100A11, ZBP1, IFITM3, PARP10 | RPS27A, RPS3, RPS14, RPL13, MT-CYB, RPS3A, RPL3, RPS4X, RPL7A, MT-CO3, EEF1A1, RPL21, RPL5, RPS8, RPS6, MT-ATP6, MT-ND4L, RPL23A, MT-ND4, MT-ND5, MT-CO2, MT-ATP8, RPL4, HOPX, EIF3L |
| **Treg** | MX1, IFI6, IFIT3, ISG15, LY6E, BST2, IFITM1, IFITM3 | RPS3A, EEF1A1, RPS4X, RPL5, RPL3, EEF2, RPL4 |

**Supplemental Table 9.** Antibodies used for flow cytometry

| Antibody | Manufacture | Clone | Cat# | Lot# | Dilution used |
|---|---|---|---|---|---|
| CD8 PerCP Cy5.5 | BD | SK1 | 341051 | 4099943 | 1:20 |
| Ki67 AF488 | BD | B56 | 561165 | 6261744 | 1:20 |
| CD38 PE | Biolegend | HB7 | 356604 | B178346 | 1:160 |
| CD38 BV421 | Biolegend | HIT2 | 303526 | B211387 | 1:20 |
| CD4 BV605 | Biolegend | RPA-T4 | 300556 | B232691 | 1:160 |
| CD19 BV785 | Biolegend | HIB19 | 302240 | B211586 | 1:160 |
| CD3 BV785 | Biolegend | OKT3 | 317330 | B231963 | 1:160 |
| IgG PE-CF594 | BD | G18-145 | 562538 | 6119766 | 1:40 |
| CD303 PeCy7 | Biolegend | 201A | 354214 | B239872 | 1:160 |
| IgA APC | Miltenyi Biotec | REA1014 | 130-116-879 | 5181030090 | 1:160 |
| CD14 AF700 | BD | M5E2 | 557923 | 7047605 | 1:80 |
| CD3 APC-Cy7 | BD | SP34-2 | 557757 | 7025869 | 1:160 |
| CD56 APC Cy-7 | Biolegend | HCD56 | 318332 | B178913 | 1:160 |
| CD56 PeCy7 | BD | B159 | 557757 | 5163892 | 1:20 |
| HLA-DR FITC | BD | G46-6 | 555811 | 28731 | 1:40 |
| CD16 PerCP-ef710 | eBioscience | CB16 | 46-0168-42 | 1991932 | 1:320 |
| IgD BV510 | Biolegend | 1A6-2 | 348220 | B226970 | 1:40 |
| IgM BV605 | BD | G20-127 | 562997 | 7026708 | 1:40 |
| CD1c BV650 | BD | F10/21A3 | 742749 | 8305935 | 1:160 |
| CD11c BUV395 | BD | B-ly6 | 563787 | 8141967 | 1:160 |
| CD27 BUV737 | BD | L128 | 564301 | 8164573 | 1:40 |
